# Reciprocal cybrids reveal how organellar genomes affect plant phenotypes

**DOI:** 10.1101/477687

**Authors:** Pádraic J. Flood, Tom P.J.M. Theeuwen, Korbinian Schneeberger, Paul Keizer, Willem Kruijer, Edouard Severing, Evangelos Kouklas, Jos A. Hageman, Raúl Wijfjes, Vanesa Calvo-Baltanas, Frank F.M. Becker, Sabine K. Schnabel, Leo Willems, Wilco Ligterink, Jeroen van Arkel, Roland Mumm, José M. Gualberto, Linda Savage, David M. Kramer, Joost J.B. Keurentjes, Fred van Eeuwijk, Maarten Koornneef, Jeremy Harbinson, Mark G.M. Aarts, Erik Wijnker

## Abstract

Assessing the impact of variation in chloroplast and mitochondrial DNA (collectively termed the plasmotype) on plant phenotypes is challenging due to the difficulty in separating their effect from nuclear derived variation (the nucleotype). Haploid inducer lines can be used as efficient plasmotype donors to generate new plasmotype-nucleotype combinations (cybrids)(Ravi et al., 2014). We generated a panel comprising all possible cybrids of seven *Arabidopsis thaliana* accessions and extensively phenotyped these lines for 1859 phenotypes under stable and fluctuating conditions. We show that natural variation in the plasmotype results in additive as well as epistatic effects across all phenotypic categories. Plasmotypes which induce more additive phenotypic changes also cause more significant epistatic effects, suggesting a possible common basis for both additive and epistatic effects. On average epistatic interactions explained twice as much of the variance in phenotypes as additive plasmotype effects. The impact of plasmotypic variation was also more pronounced under fluctuating and stressful environmental conditions. Thus, the phenotypic impact of variation in plasmotypes is the outcome of multilevel Nucleotype X Plasmotype X Environment interactions and, as such, the plasmotype is likely to serve as a reservoir of variation which is only exposed under certain conditions. The production of cybrids using haploid inducers is a quick and precise method for assessing the phenotypic effects of natural variation in organellar genomes. It will facilitate efficient screening of unique nucleotype-plasmotype combinations to both improve our understanding of natural variation in nucleotype plasmotype interactions and identify favourable combinations to improve plant performance.

## Introductory paragraph

Chloroplasts and mitochondria play essential roles in metabolism, cellular homeostasis and environmental sensing (Petrillo et al., 2014; Chan et al., 2016). Their genomes contain only a limited set of genes whose functioning requires tight coordination with the nucleus through signaling pathways that modulate nuclear and organellar gene expression (Petrillo et al., 2014; Kleine and Leister, 2016). Plasmotype variation can be strongly additive, such as in the case of chloroplast encoded herbicide tolerance (Flood et al., 2016), or can manifest in complex cytonuclear interactions as non-additive, non-linear effects (epistasis), such as found for secondary metabolites (Joseph et al., 2013). The phenotypic consequences of epistasis can be detected when a plasmotype causes phenotypic effects in combination with some, but not all nuclear backgrounds. Recent studies suggest that cytonuclear epistasis is the main route through which variation in the plasmotype is expressed (Zeyl et al., 2005; Montooth et al., 2010; Joseph et al., 2013; Joseph et al., 2013; Tang et al., 2014; Roux et al., 2016; Mossman et al., 2019) and that additive effects are both rare and of small effect.

Plasmotypic variation is relevant from an agricultural as well as evolutionary perspective (Levings, 1990; Bock et al., 2014; Dobler et al., 2014), but to understand or utilize it, it is necessary to separate nuclear from mitochondrial and chloroplastic effects. Reciprocal-cross designs, where nucleotypes segregate in different plasmotypic backgrounds, have been used to identify plasmotype-specific quantitative trait loci (Joseph et al., 2013; Tang et al., 2014), but are limited to just two plasmotypes. A larger number of plasmotypes can be studied using backcross designs where plasmotypes are introgressed into different nuclear backgrounds (Dowling et al., 2007; Sambatti et al., 2008; Miclaus et al., 2016; Roux et al., 2016), but backcross approaches are lengthy and any undetected nuclear introgressions may confound the results.

To precisely and rapidly address the contribution of organellar variation to plant phenotypes, we explored the use of a haploid inducer line available in *Arabidopsis (GFP-tailswap)* (Ravi and Chan, 2010; Ravi et al., 2014). When pollinated with a wild-type plant, the *GFP-tailswap* nuclear genome is lost from the zygote through uniparental genome elimination. This generates haploid cybrid offspring with a paternally derived nuclear genome and maternally (*GFP-tailswap*) derived mitochondria and chloroplasts (Fig. 1). These haploid plants produce stable diploid (doubled haploid) offspring following genome duplication or restitutional meiosis (Ravi and Chan, 2010). We set out to test the use of this approach to investigate how plasmotypic variation affects plant phenotypes and to what extent this variation manifests itself as additive variation or as cytonuclear epistasis.

**Figure 1.**
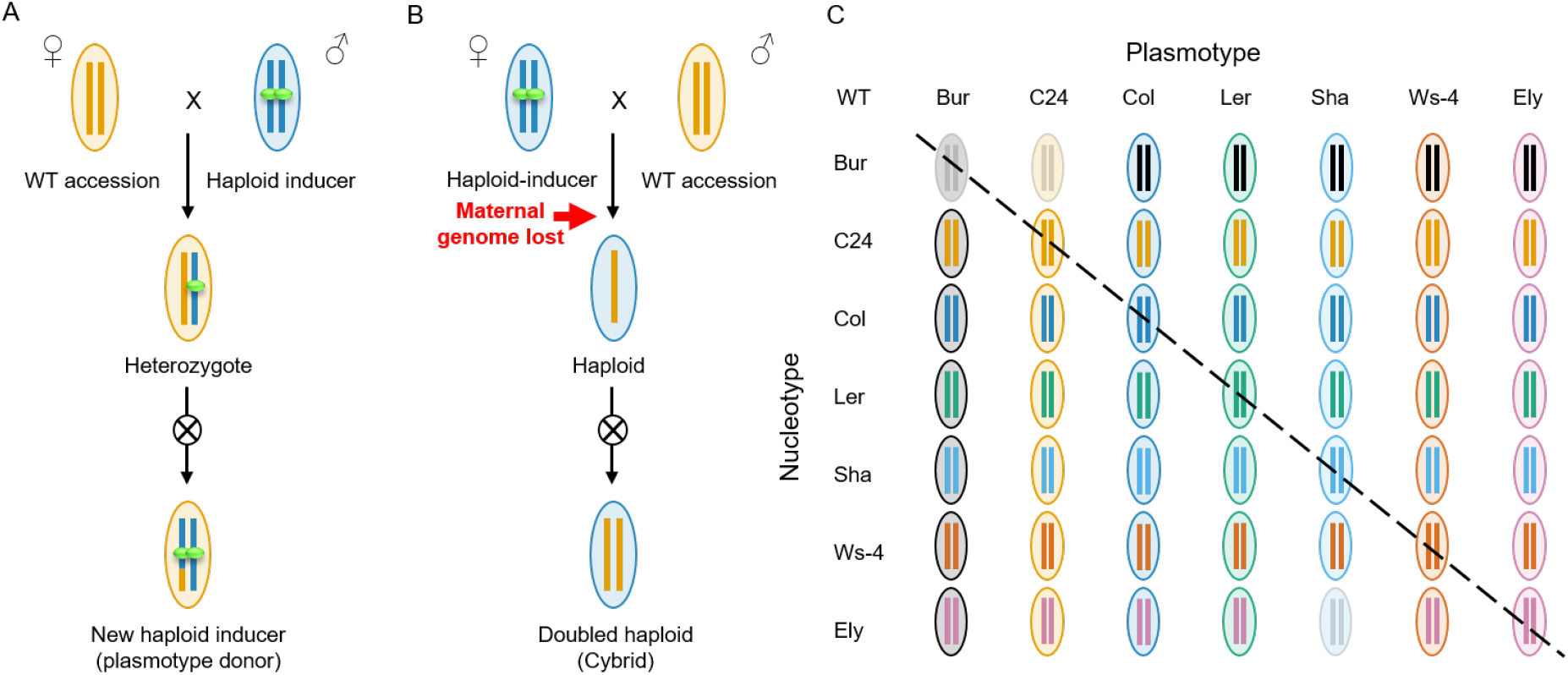
Generation of a cybrid test panel. A) Generation of a new haploid inducer (HI) line with a new plasmotype. The HI expresses a GFP-tagged *CENH3/HRT12* in a *cenh3/htr12* mutant background. A cross of a wild type (female) with a HI (male) results in a hybrid F1. A diploid F1 is selected in which no genome elimination has occurred. Self-fertilization generates an F2 population in the plasmotype of the wild-type mother. From this an F2 plant is selected that is homozygous for the *cenh3/htr12* mutation and carries the *GFP-tailswap* transgene. This F2 plant is a new HI line and can serve as plasmotype donor when used as female in crosses. Vertical bars represent the nucleotype, and the ovals represent the plasmotype. HI centromeres are indicated in green (signifying GFP-tagged CENH3/HTR12 proteins as encoded by the *GFP-tailswap* construct) that cause uniparental genome-elimination. B) HI lines can function as plasmotype donors when used as a female parent. In this case, uniparental genome elimination (red arrow) leads to a haploid offspring plant with the nucleotype of the wild-type (WT) male parent, but the plasmotype of the HI mother. C) Full diallel of all nucleotype-plasmotype combinations for which cybrids were generated. The diagonal line highlights the wild-type (WT) nucleotype-plasmotype combinations that were generated by crossing wild-type plants to plasmotype donors with the plasmotype of the wild type (self-cybrids). Bur^Bur^, Bur^C24^ and Ely^Sha^ are faded out, as they were not included in the phenotyping experiments, they have been subsequently recreated and the complete set has been submitted to NASC.

Seven different *Arabidopsis* accessions were selected for our experiment: six that represent a snapshot of natural variation (Bur, C24, Col-0, Ler-0. Sha, WS-4) and Ely, an accession with a large-effect mutation in the chloroplast-encoded *PsbA* gene (El-Lithy et al., 2005). This mutation results in reduced photosystem II efficiency (El-Lithy et al., 2005; Flood et al., 2014) and was included to evaluate the consequence of a strong plasmotype effect in our test-panel. We first generated haploid inducers for all seven plasmotypes (Fig. 1A) and then used each inducer to generate cybrid offspring for all seven nucleotypes (Fig. 1 B and C). Cybrid genotypes will henceforth be denoted as nucleotype^plasmotype^ (i.e. Ely^Bur^ denotes a cybrid with Ely nucleus and Bur plasmotype). Wild-type nucleotype-plasmotype combinations were also regenerated in this way (hereafter referred to as self-cybrids; i.e. Bur^Bur^, C24^C24^, etc.) to later compare these with their wild-type progenitors. The genotypes of all haploid cybrids were verified by resequencing. This led to the exclusion of Bur^C24^ and Bur^Bur^, because of the identification of a nucleotypic *de-novo* duplication of 200kb in these two lines, likely derived from a spontaneous duplication in a Bur wild-type progenitor used in creating these cybrids (see Online methods; Supplementary Fig. 1 to 4). With the exception of Ely^Sha^ for which we obtained seeds at a later stage, we obtained doubled haploid seeds from all haploid cybrids resulting in a testpanel of 46 cybrids and 7 wildtype progenitors. As with Ely^Sha^, Bur^C24^ and Bur^Bur^ were subsequently recreated, and the complete panel will be submitted to NASC. To visualize the genetic variation between lines within our panel we generated neighbor joining trees for the nuclear, mitochondrial and chloroplast genomes (Supplementary Fig. 5). The nucleotypes were found to be approximately equidistant, while the Ler, Ely and Col plasmotypes appear to be more closely related to each other than the other plasmotypes.

We phenotyped the cybrid panel under constant environmental conditions for absolute and relative growth rate, biomass accumulation, epinastic leaf movement, photosystem II efficiency (Φ_PSII_), non-photochemical quenching (NPQ), and elements thereof (Φ_NO_, Φ_NPQ_, q_E_ and q_I_), a reflectance-based estimate of chlorophyll, flowering time, germination, pollen abortion, and primary metabolites. To simulate more variable conditions that are frequently encountered in the field, we also screened the panel under fluctuating light for all the above-mentioned photosynthesis-related phenotypes and assayed germination rates under osmotic stress and after a controlled deterioration treatment. Counting individual metabolite concentrations and single time points in the time series separately, we collected in total 1859 phenotypes (Supplementary Data 1, Supplementary Table 4). To avoid overrepresentation of highly correlated and non-informative phenotypes we selected a subset of 92 phenotypes (Online methods, Supplementary Table 2) comprising 24 from constant growth conditions, 32 from fluctuating or challenging environmental conditions and 36 primary metabolites for further analysis (Supplementary Fig. 6, Supplementary Table 2).

Comparison of six self-cybrids with their genetically identical wild-type progenitors for these 92 phenotypes did not reveal significant phenotypic differences (Supplementary Table 1) from which we infer that uniparental genome elimination is a robust method to generate cybrids. To determine the relative contributions of nucleotype, plasmotype, and their interaction to the observed phenotypic variation, we estimated the fraction of the broad sense heritability (H^2^; also called repeatability (Falconer and Mackay, 1996)) explained by each. Across the entire panel the average contribution to H^2^ of nucleotype, plasmotype and nucleotype-plasmotype interaction was 65.9%, 28.0% and 6.1% respectively (Supplementary Table 2 and 3; Supplementary Data 2). Most of the plasmotype derived additive variation was caused by the Ely plasmotype, arising from the *psbA* mutation. When this plasmotype was excluded from the analysis, the nucleotype, plasmotype and their interaction account for 91.9%, 2.9% and 5.2% of the genetic variation, respectively (Supplementary Table 2 and 3; Supplementary Data 2). So, while nucleotype-derived additive variation is the main genetic determinant of the cybrid phenotype, variation caused by plasmotype additive effects as well as epistatic effects results in substantial phenotypic differences.

Next we sought to assess whether there are general patterns in how specific nucleotypes and plasmotypes interact. To this end we first assessed which plasmotype changes result in additive phenotypic changes. Plasmotype replacements involving the Ely plasmotype lead to additive changes in, on average, 50 (out of 92) phenotypes across the 7 nucleotypes (Table 1A). Changes involving the Bur plasmotype lead to on average 10 significant additive effects, 8 of which are photosynthesis-related (Supplementary Data 2). Other plasmotype changes show on average one additive effect, in predominantly non-photosynthetic phenotypes. Comparison of wild-type cytonuclear combinations with all their iso-nuclear cybrid lines also shows that plasmotype changes involving Ely and Bur plasmotypes show the most epistatic effects (on average 43 and 6 respectively) (Table 1B). The number of epistatic effects resulting from the Bur plasmotype range between 0 (L*er*^L*er*^ vs L*er*^Bur^) to 10 (Sha^Sha^ vs Sha^Bur^), indicating high variability. Plasmotype changes involving other plasmotypes show more modest numbers of significant epistatic effects that range from 0 to 6. Plasmotypes that result in more additive effects also cause more epistatic effects (Pearson correlation coefficient of 0.99, p-value 1.3e-5) suggesting a possible common cause (Supplementary Fig. 7).

**Table 1.** Significant plasmotype induced effects in 92 phenotypes. A) Number of observed significant plasmotype additive effects when a specific plasmotype is changed for another plasmotype, regardless of the nucleotype. Note that the replacement of Bur (top row) and Ely plasmotypes (last column) result in most plasmotype additive effects. B) Number of observed significant epistatic effects in phenotypes between wild-type nucleotype-plasmotype combinations and cybrids with different plasmotypes. Rows indicate the number of significant effects when comparing self-cybrids to cybrids with identical nucleotype but non-native plasmotype. Columns indicate specific plasmotype changes. Note that changing the Ely plasmotype for another plasmotype (bottom row and last column) results in many epistatic effects due to the large-effect mutation in the chloroplast-encoded *PsbA* gene of the Ely plasmotype. Similar effects, but of smaller magnitude, result from changing the Bur plasmotype (top row and first column). Posthoc tests were used with Hochberg’s p-value correction for panel A and Dunnett’s p-value correction (with the wild-type as control) for panel B, α = 0.05. nd = not determined. For underlying p-values and phenotypes see Supplementary Data 2. Yellow cells indicate low number of significant effects; blue cells show higher number of significant effects.

| # of significant phenotypes<br>0 55 |  | Plasmotype |  |  |  |  |  |  |
| --- | --- | --- | --- | --- | --- | --- | --- | --- |
|  |  | XXX<sup>Bur</sup> | XXX<sup>C24</sup> | XXX<sup>Col</sup> | XXX<sup>Ler</sup> | XXX<sup>Sha</sup> | XXX<sup>Ws-4</sup> | XXX<sup>Ely</sup> |
| Plasmotype | XXX<sup>Bur</sup> |  | 12 | 15 | 10 | 15 | 6 | 55 |
|  | XXX<sup>C24</sup> |  |  | 1 | 0 | 1 | 0 | 50 |
|  | XXX<sup>Col</sup> |  |  |  | 2 | 2 | 1 | 50 |
|  | XXX<sup>Ler</sup> |  |  |  |  | 0 | 1 | 48 |
|  | XXX<sup>Sha</sup> |  |  |  |  |  | 2 | 49 |
|  | XXX<sup>Ws-4</sup> |  |  |  |  |  |  | 49 |
|  | XXX<sup>Ely</sup> |  |  |  |  |  |  |  |

| # of significant phenotypes<br>0 48 |  | Plasmotype |  |  |  |  |  |  |
| --- | --- | --- | --- | --- | --- | --- | --- | --- |
|  |  | XXX<sup>Bur</sup> | XXX<sup>C24</sup> | XXX<sup>Col</sup> | XXX<sup>Ler</sup> | XXX<sup>Sha</sup> | XXX<sup>Ws-4</sup> | XXX<sup>Ely</sup> |
| wildtype nucleotype-plasmotype combination | Bur wildtype |  | nd | 4 | 7 | 9 | 4 | 48 |
|  | C24<sup>C24</sup> | 4 |  | 1 | 0 | 3 | 1 | 32 |
|  | Col<sup>Col</sup> | 5 | 2 |  | 0 | 1 | 1 | 39 |
|  | Ler<sup>Ler</sup> | 0 | 0 | 1 |  | 3 | 6 | 37 |
|  | Sha<sup>Sha</sup> | 10 | 2 | 1 | 1 |  | 2 | 40 |
|  | Ws-4<sup>Ws-4</sup> | 4 | 3 | 0 | 0 | 4 |  | 37 |
|  | Ely<sup>Ely</sup> | 41 | 45 | 44 | 42 | nd | 42 |  |

Though the average total explained variance due to the cytonuclear epistasis is only 5.2%, these interactions can have strong effects for specific phenotypes or in specific cybrids. Explained variance for some phenotypes can be markedly higher, for example for projected leaf area this amounts to 12.3%, for hyponastic leaf movement to 8.3% and for ΦNPQ to 17.8%. A strong epistatic effect in pollen abortion (43.5%) was due to relatively high pollen abortion in Sha^Sha^ (Fig. 2A) that we also observed in Sha wildtype. The increased pollen abortion in its native nucleotype is surprising and could indicate either incomplete compensation due to the accumulation of deleterious variants or perhaps to facilitate increased outcrossing. The only cybrid for which we initially failed to obtain seed was Ely^Sha^. This haploid was regenerated and pollinated with wild-type Ely pollen to increase the chance of seed set. The diploid offspring showed 45% of pollen abortion and were male sterile, indicating that in combination with the Ely nucleotype the Sha plasmotype results in cytoplasmic male sterility (Supplementary Fig. 9). In combination with the Sha plasmotype pollen abortion across the seven nucleotypes can range from near zero, to 8.9% in Sha^Sha^ and to full male sterility in Ely^Sha^, highlighting the strong epistasis that can be present.

**Figure 2.**
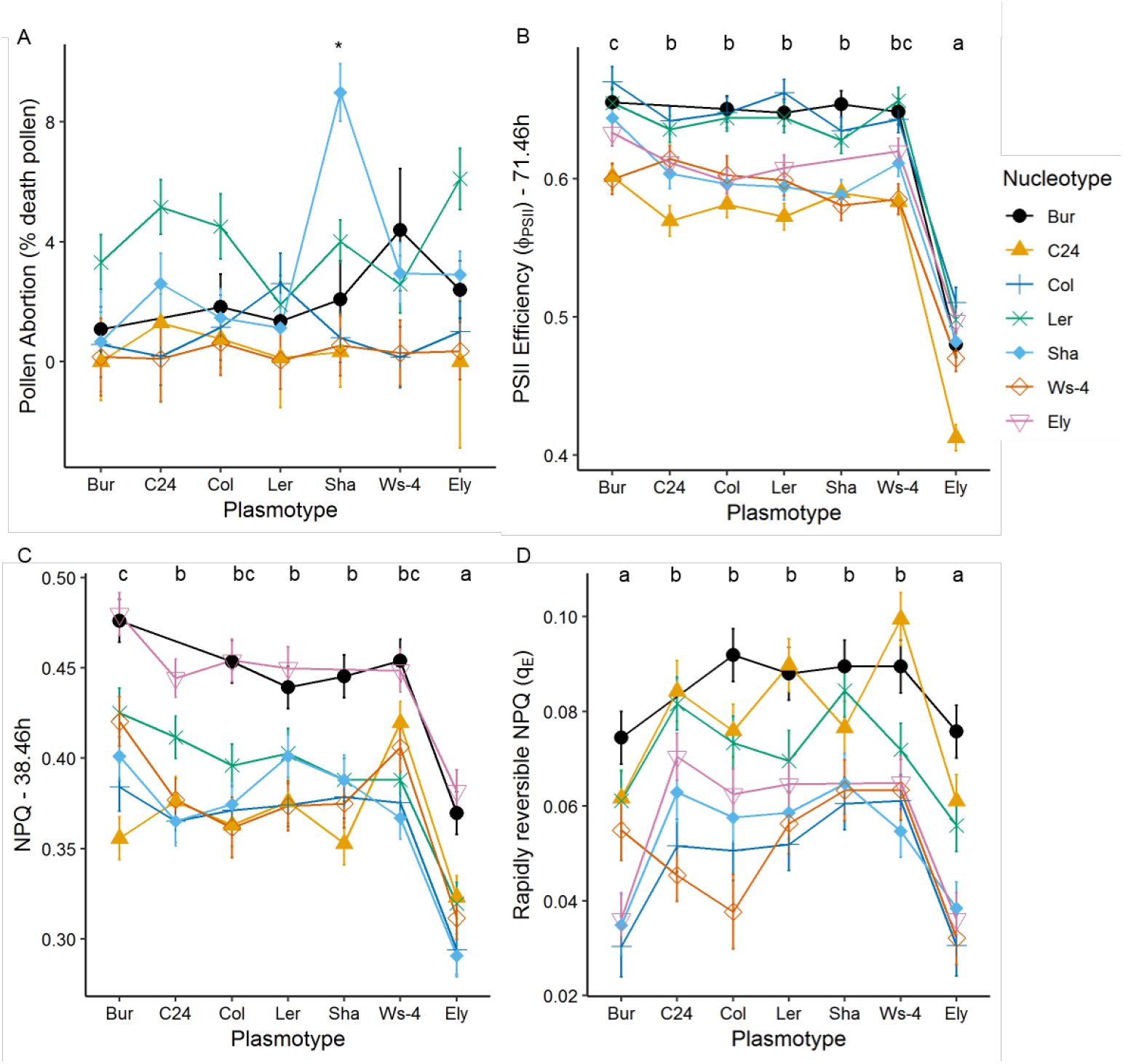
Plasmotype changes result in cytonuclear epistasis, and in the case of cybrids with the Ely and Bur plasmotype also in additive effects. A) Pollen abortion, percentage of dead pollen out of 250. B) PSII efficiency (Φ_PSII_) 71.46 hours after start of experiment, after a full day of fluctuating light with a maximum difference between 500 and 100 μmol/m^2^/s irradiance (see Fig. 3C for light treatment). C) NPQ at 38.46 hours after start of experiment, which is at 300 μmol/m^2^/s on a sigmoidal light curve starting at 65 μmol/m^2^/s. D) The rapidly reversible component of NPQ, q_E_, at 259 μmol/m^2^/s after a full day of fluctuating light with a maximum difference between 500 and 100μmol/m^2^/s. X-axis are labelled with the plasmotype, and the colours represent the nucleotypes. Any deviation from a horizontal line represents a potential additive or epistatic effect. Error bars represent the standard error of the mean. The * in panel A indicates a unique significant difference between the Sha^Sha^ cybrid and other cybrids with Sha nucleotypes (epistasis) (Hochberg’s test, n=4-10). The letters above panels B, C and D represent significant differences between plasmotypes regardless of the nucleotype (additivity) (Hochberg’s test, n=4*7). For panels B, C and D plants were grown at 200 μmol/m^2^/s for 21 days prior to starting the experiment.

Cybrids with the Ely plasmotype exhibit clear additive effects: all have a lower PSII efficiency (Φ_PSII_) (Fig. 2B) and lower values for other photosynthesis related phenotypes i.e. NPQ, q_E_ and chlorophyll content (Fig. 2C and Supplementary Data 2). This reduced Φ_PSII_ is likely to be responsible for the concomitant reductions in biomass, growth rate and seed size and altered primary metabolite content (Supplementary Data 2). To test whether additive effects could also be detected at the level of gene expression we contrasted the transcriptome of Ely^Ely^ with that of the Ely^L*er*^ and Ely^Bur^ cybrids. We also compared the transcriptomes of L*er*^L*er*^, L*er*^Bur^, and L*er*^Ely^ (Supplementary Data 3; for details see Supplementary Fig. 10 and Supplementary Table 5). Exchanging the Ely plasmotype with Ler or Bur, in either the Ler or Ely nuclear background resulted in a consistent change in the expression of 40 genes, of which most were upregulated (Supplementary Table 6). A GO-term analysis revealed that these genes are significantly enriched for those involved in photorespiration (GO:0009853) and in glycine- and serine family amino acid metabolism (GO:0006544 and GO:0009069) (Supplementary Data 3). This is in line with the low serine and glycine content of cybrids with the Ely plasmotype which suggests reduced photorespiration (Supplementary Data 2) (Somerville and Ogren, 1980) and can be linked to lower overall photosynthetic activity.

The Ely plasmotype was deliberately included in our panel for its strong additive effect. In addition to Ely we also observed strong additive effects from the Bur plasmotype which are mainly restricted to the photosynthetic parameters. Under normal conditions PSII efficiency is slightly increased by the Bur plasmotype (1.6%), however when fluctuating the light intensity, this difference becomes more apparent (3.5% increase) (Fig. 2B and 3). This increase in Φ_PSII_, under fluctuating conditions results in a corresponding reduction in Φ_NO_ and Φ_NPQ_ of 7.3% and 2.2% respectively. NPQ, q_E_ and q_I_ are also influenced by the plasmotype, but the time points at which these differences occur differs per phenotype (Fig. 3A and B). The Bur plasmotype increases NPQ, with the largest increase of 5.9% at the beginning of day 2 (38.46h) (Fig. 2C), while the rapidly reversible component of NPQ, qE, has a maximum reduction of 26.6% at the end of day 3 (71.46h) (Fig. 2D).

**Figure 3.**
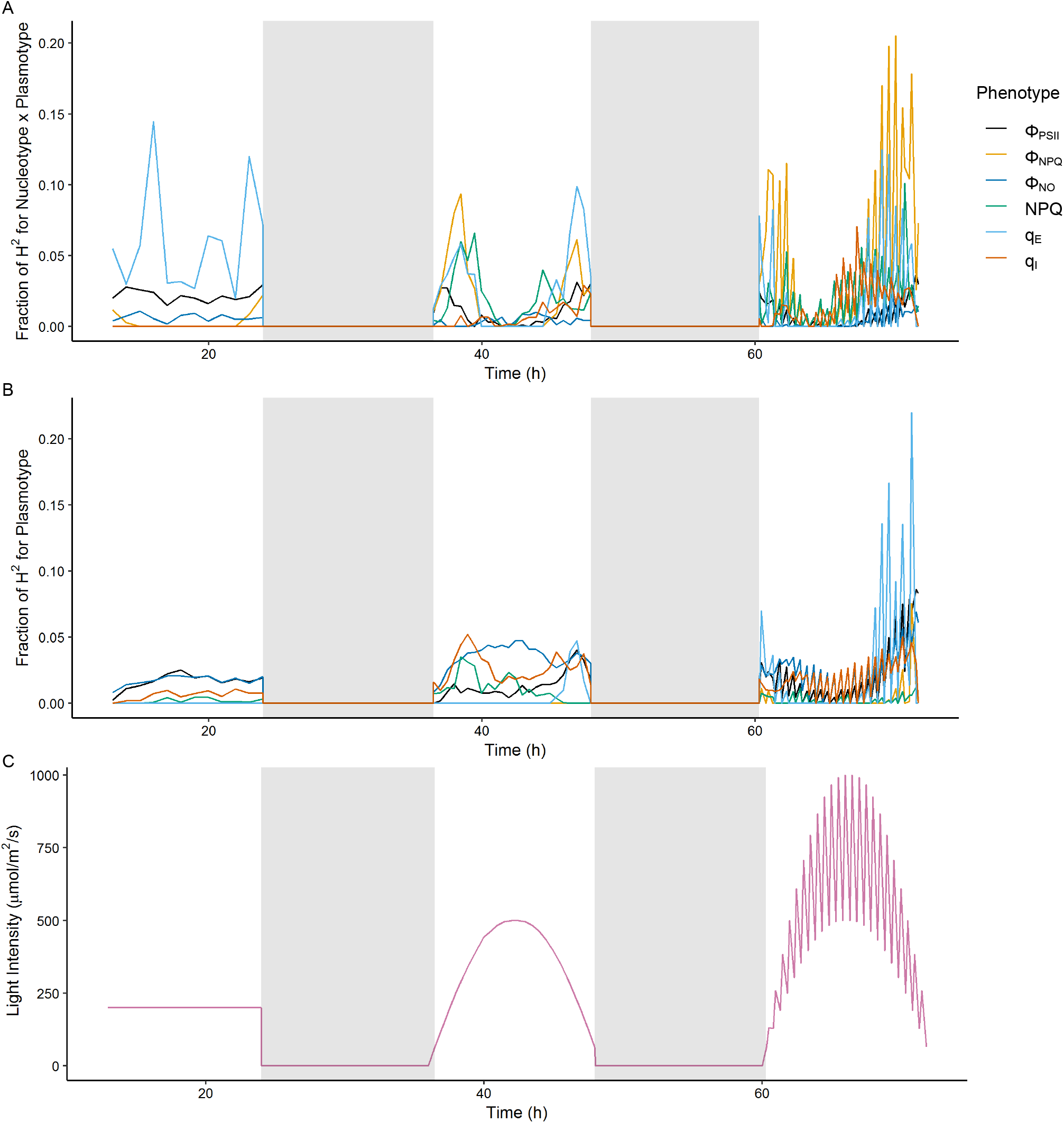
The fraction of explained genetic variation (H^2^) for photosynthesis phenotypes differs depending on light conditions. A) shows the fraction of H^2^ for plasmotype epistatic effects. B) shows the fraction of H^2^ for plasmotype additive effects. C) shows the light intensity for three consecutive days with growth under steady light (day 1), sinusoidal light intensity (day 2) and fluctuating light intensity (day 3). Days are separated by nights (shaded areas). Note that the fraction of H^2^ for different phenotypes changes markedly during days 2 and 3. Some phenotypes are explained largely by additive effects (i.e. q_E_) while others by interaction (i.e. Φ_NPQ_). A replication of this experiment is shown in Supplementary Fig. 8.

These photosynthesis-related phenotypes are likely to be due to chloroplast-derived variation. In support of a chloroplastic origin for this photosynthetic variation, measurements of mitochondrial respiration suggest that Bur is not an outlier and shows standard respiration rates (Supplementary Fig. 11). Based on coverage plots there are no obvious duplications or deletions in the mitochondrial or chloroplast sequence of Bur thus we expect that altered expression or protein activity as opposed to gene gain or loss is driving the Bur derived phenotypes (Supplementary Fig. 12). We annotated the sequence variation of all plasmotypes using SnpEff (Cingolani et al., 2012). From this we found no large effect mutations in the Bur mitochondria. There were, however, unique missense variants in the chloroplastic genes MATURASE K (MATK), NAD(P)H-QUINONE OXIDOREDUCTASE SUBUNIT 6 (NDHG) and chloroplast open reading frame 1 (YCF1) as well as a frameshift mutation in tRNA-Lys (TRNK) (Supplementary Data 4). NDHG is part of the NAD(P)H-dehydrogenase-like complex (NDH). NDH is located inside the thylakoid membrane and acts, amongst others, as a proton pump in cyclic electron flow around photosystem I and chlororespiration. NDH creates a pH differential that can be causative of the observed non-photochemical quenching phenotypes (Strand et al., 2017; Laughlin et al., 2019). In contrast to Ely, the plasmotype which evolved in response to the use of herbicides, an anthropogenic selective pressure (Flood et al., 2016), the Bur plasmotype represents a naturally occurring plasmotype that has an additive impact on key photosynthetic phenotypes.

Our experiments have shown that a clean, systematic exploration of plasmotypic variation in a plant species is feasible. To our knowledge, apart from the *cenh3* mutant used here, there is only one other intraspecific haploid inducer available (the maize *ig* mutant) which can be used via the maternal line and thus replace the plasmotype (Kermicle, 1969; Schneerman et al., 2000; Houben et al., 2011). Current knowledge of *cenh3* mediated uniparental genome elimination should allow for the creation of maternal haploid inducers in a wider range of species (Karimi-Ashtiyani et al., 2015). This would allow elite nucleotypes to be brought into new plasmotypic backgrounds to explore novel plasmotype-nucleotype combinations. Our data indicate that there is substantial variation for phenotypes such as NPQ and Φ_PSII_ which are important for plant productivity (Flood et al., 2011; Kromdijk et al., 2016; Murchie et al., 2018). Next to Ely, we identified one new plasmotype (Bur) that significantly impacts photosynthesis in an additive manner. Expanding our panel would likely find more, suggesting that future research aiming to enhance crop photosynthesis should play close attention to available plasmotypic variation. Apart from studying natural variation, the use of haploid inducers as plasmotype donors could be used to transfer cytoplasmic male sterility (CMS), herbicide resistances or genetically engineered plasmotypes. Plant plasmotypes, are notoriously difficult to genetically modify, although recently there have been some advances in this regard (Jin and Daniell, 2015; Zhang et al., 2015; Kwak et al., 2019; Ruf et al., 2019). The use of haploid inducers as plasmotype donors could further increase the accessibility of such modifications as transformations could be undertaken in a compatible nucleotype and once achieved can be transferred into different nucleotypes, thus amplifying the potential impact of successful plasmotype modifications.

Exploring the potential of plasmotypic variation via the use of haploid inducer lines is not only promising for plant breeding, but also for understanding the role such variation plays in plant adaptation (Bock et al., 2014; Dobler et al., 2014). Our results show that despite considerable genetic divergence between the genotypes used in our panel, all cybrids were viable, this in itself suggests a remarkable degree of conservation for the fundamental components of cytonuclear interactions. Although we do find clear additive effects of some plasmotypes, the majority of the plasmotype derived variation manifests as epistasis in the traits we measured which is in line with previous research in plants, animals, and fungi (Zeyl et al., 2005; Dowling et al., 2007; Montooth et al., 2010; Joseph et al., 2013; Roux et al., 2016). Also in line with studies of mitonuclear interactions in animals is the observation that phenotypic variation due to plasmotypic variation becomes more pronounced under fluctuating and stressful conditions (Dowling et al., 2007; Hoekstra et al., 2013; Mossman et al., 2016; Hill et al., 2019). Both our results and previous work suggests that multilevel interactions (i.e. Nucleotype x Plasmotype x Environment) may be the primary mechanism by which plasmotypic variation is expressed. Thus, plasmotypic variation may act as an evolutionary capacitor providing novel phenotypes in specific genetic and environmental contexts, such variation may be particularly important for both crops and wild species in our rapidly changing climate.

## Online methods

### Plant materials

Seven *Arabidopsis* accessions were chosen for the construction of a full nucleotype-plasmotype diallel. Ely (CS28631) is atrazine resistant due to a chloroplast-encoded mutation in *PsbA* which leads to a modified D2 protein that greatly reduces PSII efficiency (El-Lithy et al., 2005). Ws-4 (CS5390) was included for its unusual photosystem II phosphorylation dynamics (Yin et al., 2012). Bur (CS76105) is commonly used in diversity panels and is a standard reference accession. Sha (CS76227) was selected based on its capacity to induce cytoplasmic male sterility in some crosses (Gobron et al., 2013). The set was completed by adding Ler (CS76164), Col (CS76113) and C24 (CS76106) which are three widely used genotypes in *Arabidopsis* research. Col is the reference genome for nuclear and chloroplast sequences and C24 for the mitochondrial sequence. The *GFP-tailswap* haploid-inducer that expresses a GFP-tagged CENTROMERE HISTONE 3 protein in a *cenh3/htr12* mutant background, is in a Col background (Ravi and Chan, 2010).

### Generation of a nucleotype-plasmotype diallel

To generate new nucleotype-plasmotype combinations, plants of all seven accessions (Bur, C24, Col, Ely, Ler, Sha and Ws-4) were crossed as males to *GFP-tailswap* resulting in all cybrids with the Col plasmotype. New HI lines were created by crossing the original *GFP-tailswap* line as a male to the six additional plasmotype mothers (Bur, C24, Ely, L*er*, Sha and Ws-4). Genome elimination does not always occur and some of the offspring were diploid F1 lines. These were selfed and F2 lines homozygous for the *cenh3/htr12* mutation and carrying the *GFP-tailswap* were selected as new HI lines in different plasmotypic backgrounds (Fig. 1A). Plants of all seven accessions were then crossed as males to these new HI lines and the haploids arising from these 49 crosses were identified based on their phenotype (as described in Wijnker et al. (2014)). These haploid lines self-fertilized, either following somatic genome duplication or after restitutional meiosis (Ravi and Chan, 2010), and gave rise to doubled haploid offspring (Fig. 1B). The resulting 49 lines comprise a full diallel of 21 pairs of reciprocal nucleotype-plasmotype combinations (cybrids) as well as seven nucleotype-plasmotype combinations that have, in principle, the same nucleotype-plasmotype combinations as their wild-type progenitors (self-cybrids; Fig. 1C, diagonal). All cybrids and the wild-type accessions were propagated for one generation before use in further experiments, with the exception of Ely^Sha^ of which the original haploid died without setting seed and was recreated at a later stage by generating haploids that were pollinated with Ely wild-type plants to ensure seed set.

### Genotype confirmation

To confirm that all cybrids in our panel are authentic, all 49 cybrids and 7 wild-type progenitors were whole-genome sequenced at the Max Planck Genome Centre Cologne (Germany) using Illumina Hiseq 2500 150-bp paired-end sequencing. The cybrids were sequenced at 8.5X coverage and the wild-type progenitors at 40X coverage. To remove erroneous bases, we performed adapter and quality trimming using Cutadapt (version 1.18) (Martin, 2011). Sequences were clipped if they matched at least 90% of the total length of one of the adapter sequences provided in the NEBNext Multiplex Oligos for Illumina^®^ (Index Primers Set 1) instruction manual. In addition, we trimmed bases from the 5’ and 3’ ends of reads if they had a phred score of 20 or lower. Reads that were shorter than 70 bp after trimming were discarded. Trimmed reads were aligned to a modified version of the A. thaliana Col-0 reference genome (TAIR10, European Nucleotide Accession number: GCA_000001735.2) which contains an improved assembly of the mitochondrial sequence (Genbank accession number: BK010421) (Sloan et al., 2018) using bwa mem (version 0.7.10-r789) (Li, 2013) with default parameters. The resulting alignment files were sorted and indexed using samtools (version 1.3.1) (Li et al., 2009). Duplicate read pairs were marked using the MarkDuplicates tool of the GATK suite (version 4.0.2.1), using an optical duplicate pixel distance of 100, as recommended in the documentation of GATK when working with data from unpatterned Illumina flowcells. Variants were called using a workflow based on GATK Best Practices. Base quality scores of aligned reads were recalibrated using GATK BaseRecalibrator with default parameters, using a set of variants of a world-wide panel of 1135 *A. thaliana* accessions (The 1001 Genomes Consortium, 2016) (obtained from ftp://ftp.ensemblgenomes.org/pub/plants/release-37/vcf/arabidopsis_thaliana/) as known sites. Following base recalibration, variants were called in each sample using GATK HaplotypeCaller, allowing for a maximum of three alternate alleles at each site. Samples were then jointly genotyped using GATK GenomicsDBImport and GATK GenotypeGVCFs with default parameters. This last step generated three different VCF files: one containing the calls of the nuclear genome, one containing calls of the mitochondrial genome and one containing calls of the chloroplast genome.

To remove likely false positive calls, we filtered the callsets using two complementary approaches. First, we filtered the nuclear callset using GATK VariantRecalibrator and GATK ApplyVQSR (--truth-sensitivity-filter-level set at 99.9), using the set of variants called in the world-wide panel of 1135 A. thaliana accessions as a training and truth set (prior=10.0). This step could not be performed for the mitochondrial and chloroplast calls, as these lack a golden truth set that can be used for recalibration. Second, we filtered variants based on their quality by depth score (QD). For the nuclear callset, we used a QD score of 40, leaving 3.7 million SNPs, for the chloroplast callset a QD of 25, leaving 356 SNPs and for the mitochondrial callset a QD of 20, leaving 135 SNPs.

46 cybrids were found to have the correct genotypes. With one line, Bur^Ws-4^, there was a sample mix-up during library preparation with Sha^Sha^. Leading to two Sha^Sha^ samples and no sequenced Bur^Ws-4^ sample. Fortunately, we did have a true Bur^Ws-4^ cybrid, which we confirmed via both phenotype (Bur and Sha nucleotypes are phenotypically distinct from one another) and genotype through the KASP^TM^ markers (see below) (Supplementary Table 8). To confirm the Sha cybrids we therefore used the Sha genotype (CS76382) from the 1001 genomes project (The 1001 Genomes Consortium, 2016). Two other lines, C24^C24^ and Ws-4^Col^, had a high number of heterozygous calls in their plasmotypes, with C24^C24^ being heterozygous with C24^Col^ and Ws-4^Col^ being heterozygous with Ws-4^Bur^. As in the production of Ws-4^Col^ no plant was used with a Bur plasmotype, this cannot be heteroplasmy. To ensure that this was a sample mix-up and the putative event of cross-contamination had occurred in the laboratory, we designed KASP^TM^ makers (LGC, https://www.lgcgroup.com) and genotyped all lines. These KASP^TM^ markers are designed to be unique to the chloroplast of one accession, and designed on SNPs that were called heterozygous in the sequence analysis (Supplementary Table 7). The KASP^TM^ assay can distinguish between both homozygouse or heterozygous states. We ran all seven KASP^TM^ markers on all lines, for C24^C24^ and Ws-4^Col^ this included plants from the same seed batch as the plants used for sequencing, as well as direct offspring of the sequenced plants. All lines showed the correct genotypes, and no heterozygosity was observed in any of the lines, including C24^C24^ and Ws-4^Col^ (Supplementary Table 8). Unfortunately, the Ely^Sha^ used for sequencing died before setting seed and although it has since been recreated, it could not be included in our phenotypic analyses. We have used the KASP^TM^ marker for the Sha chloroplast, and confirmed it to be correct (Supplementary Table 8).

To check for any incomplete chromosome elimination, we calculated the read coverage for all cybrids, normalized per chromosome. We did not observe any remaining chromosomes, although we found a 200kb duplication of nuclear DNA in Bur^Bur^ and Bur^C24^. In Bur^C24^ and the self-cybrid Bur^Bur^ we discovered the presence of a duplicated segment on chromosome 2. Because this duplicated segment is present (and identical) in two independent cybrid lines and this segment is of a Bur nuclear origin (i.e. there are only Bur SNPs in this region), we conclude this segment results from a *de-novo* duplication in one of the wild-type Bur lines used to generate these cybrids. Following the exclusion of phenotyping data for Bur^Bur^ and Bur^C24^ we limited our analyses to 46 rather than 49 cybrids. The parental lines were included in the screens to test for possible unforeseen effects of cybrid production (which involves a haploid growth stage). This brings the number of phenotyped lines in this study to a total of 53 (40 cybrids, 6 self-cybrids and 7 wild types).

The fuctional effects of the chlorplastic and mitochondrial SNPs and INDELs were predicted using SnpEff (Cingolani et al., 2012). A SnpEff database was built using the genome, transcriptome and proteome as released in TAIR10.1. SNPs and INDELs were predicted on the filtered VCF, as mentioned above. In the analysis we only considered varaints with a “HIGH” or “MODERATE” impact.

### Phenotyping

Cybrids were phenotypically assessed using different platforms. For details on the number of phenotypes per experiment see Supplementary Table 4.

Growth, PSII efficiency (Φ_PSII_), chlorophyll reflectance and leaf movement (all parameters at n=24) was screened in the Phenovator platform, a high-throughput phenotyping facility located in a climate-controlled growth chamber (Flood et al., 2016). This phenotyping platform measured the plants for: Φ_PSII_ using chlorophyll fluorescence, reflectance at 480 nm, 532 nm, 550 nm, 570 nm, 660 nm, 700 nm, 750 nm and 790 nm, and projected leaf area (PLA) based on pixel counts of near infra-red (NIR) images (Flood et al., 2016). The growth chamber was set to a 10 h day/14 h night regime, at 20°C day and 18°C night temperature, 200 μmol m^-2^ s^-1^ irradiance, and 70% relative humidity. The plants were grown on a rockwool substrate and irrigated daily with a nutrient solution as described in Flood et al. (2016).

Growth (n=24) and subsequently above ground biomass (n=12) was measured in another high-throughput phenotyping facility (Kokorian et al., 2010), where projected leaf area was measured three times per day with 14 fixed cameras (uEye Camera, IDS Imaging Development Systems GmbH, Obersulm, Germany). This growth chamber was set to a 10 h day/14 h night regime, at 20°C day and 14°C night temperature, 200 μmol m^-2^ s^-1^ light and 70% relative humidity. Plants were grown on rockwool and irrigated weekly with a nutrient solution as described before.

Non-fluctuating and fluctuating light treatments were performed in the DEPI phenotyping facility of Michigan State University (n=4)(Cruz et al., 2016). This facility is able to measure the chlorophyll fluorescence derived photosynthetic parameters, Φ_PSII_, Φ_NO_, Φ_NPQ_, NPQ, q_E_, q_I_. Three-week-old plants were moved into the facility, where they were left to acclimatize for 24 hours after which three days of phenotyping was performed under different light regimes. On the first day the plants were illuminated with a constant light intensity of 200 μmol m^-2^ s^-1^. On the second day the plants received a sinusoidal light treatment where the light intensity began low and gradually increased to a maximum of 500 μmol m^-2^ s^-1^ light from which it deceased back down to 0. On the third day the plants received a fluctuating light treatment ranging between 0 and 1000 μmol m^-2^ s^-1^ light in short intervals (Fig. 3C). For the second experiment in the DEPI phenotyping facility the experiment was extent with 2 days, in which day 4 replicated day 2 and day 5 replicated day 2 (Supplementary Data 1 and Supplementary Fig. 8C). For further details see Cruz et al. (2016).

Bolting time and flowering time were measured on all cybrids (n=10) in a greenhouse experiment in April 2017, with the exception of Ely nucleotype cybrids which needed vernalisation and were not included in this experiment. Additional lighting was turned on when the natural light intensity fell below 685.5 μmol m^-2^ s^-1^, and turned off when the light intensity reached 1142.5 μmol m^-2^ s^-1^, with a maximum of 16 h per day.

Seeds for the germination experiments were generated from two rounds of propagation. In the first-round seeds were first sown in a growth chamber set to a 10 h day/14 h night regime, at 20°C day and 18°C night temperature. 200 μmolm^-2^s^-1^ light intensity, and 70% relative humidity. After three weeks they were moved to an illuminated cold room at 4°C for six weeks of vernalization. After vernalization all plants (n=8) were moved to a temperature-controlled greenhouse (20°C) for flowering and seed ripening. Exceptions to this were L*er*^Ely^, L*er*^Ws-4^, and Ely^Ws-4^ for which no doubled haploid seed was available at the beginning of the first propagation round. L*er*^Wly^ and L*er*^Ws-4^ were sown later, during the vernalization stage and flowered at the same time as the vernalized plants. Ely^Ws-4^ produced haploid seed at a later stage and could not be included in the first propagation round. Plants were grown in a temperature-controlled greenhouse set at 20°C. In this round only lines with the Ely nucleotype were vernalized. For the germination experiments seeds were stratified on wet filter paper for four days at 4°C before being assayed in the Germinator platform (Joosen et al., 2010) for seed size, germination rate and total germination percentage. Germination under osmotic stress was performed on filter paper with 125 mM NaCl. For the controlled deterioration treatment, seeds were incubated for 2.5, 5 or 7 days at 40°C and 82% RH and subsequently assayed in the Germinator platform without stratification.

To assess pollen abortion all cybrid lines and wild-type progenitors (except those with the Ely nucleotype) were grown simultaneously in a growth chamber (Percival) under controlled conditions (16H/ 8H light cycle, 21°/18° °C and 50%-60% relative humidity). Pollen abortion was manually assessed for all the ecotypes by using a differential staining of aborted and non-aborted pollen grains (Peterson et al., 2010). A total of three plants and three flowers per plant of each cybrid were collected on the same day and submerged in a drop of 13 ul of phenol-free Alexander staining solution placed on a glass slide with a glass cover slip of 18×18 mm. For each flower 250 pollen grains were counted and the number of aborted pollen therein.

Oxygen consumption of seedlings was measured in 2 mL of deionized water with a liquid-phase Oxytherm oxygen electrode system (Hansatech Instruments) calibrated at the measurement temperature. Three-day-old seedlings (about 50 mg) were directly imbibed in the electrode chamber. The rates of oxygen consumption were measured after tissue addition and subtracted from the rates after addition of 500 μM KCN. Results are the mean of at least five measurements. Measurements for different genotypes were performed on consecutive days, and to correct for daily variation, normalized to Col-0 samples that were run daily.

### Metabolomics

Plant material for primary metabolite analysis was obtained from the ‘Phenovator’ photosynthetic phenotyping experiment. Plants were harvested 26 days after sowing, which due to the 10-hr photoperiod was prior to bolting for all lines. Samples were frozen in liquid nitrogen, and samples of each genotype were subsequently combined into four pools each made up of material of approximately six replicates. Each pool was ground and homogenized before an aliquot was taken for further analysis. Reference samples for the metabolite analysis were composed of material from all seven parents in equal amounts and then homogenized. The method used for the extraction of polar metabolites from *Arabidopsis* leaves was adapted from Lisec et al. (2006) as described by Carreno-Quintero et al. (2012). Specific adjustments for *Arabidopsis* samples were made as follows; the polar metabolite fractions were extracted from 100 mg of *Arabidopsis* leaf material (fresh weight, with max. 5% deviation). After the extraction procedure, 100 μL aliquots of the polar phase were dried by vacuum centrifugation for 16 hours. The derivatization was performed on-line similar as described by Lisec et al. (2006) and the derivatized samples were analyzed by a GC-ToF-MS system composed of an Optic 3 high-performance injector (ATAS^TM^, GL Sciences, Eindhoven, The Netherlands) and an Agilent 6890 gas chromatograph (Agilent Technologies, Santa Clara, California, United States) coupled to a Pegasus III time-of-flight mass spectrometer (Leco Instruments, St. Joseph, Michigan, United States). Two microliters of each sample were introduced in the injector at 70°C using 5% of the sample (split 20). The detector voltage was set to 1750 Volts. All samples were analyzed in random order in four separate batches. The systematic variation that inadvertently is introduced by working in batches, was removed upon analysis of covariance. In this model the batch number was used as a factor (four levels) and “run number within a batch” as a covariate since it is also expected that (some) variation will be introduced by the sample run order within each batch. For this the S2 method described by (Wehrens et al., 2016) was used to perform the least-squares regression. After quality control and removing metabolites with more than 20% missing data and a broad sense heritability (H^2^) of less than 5%, we were left with data on 41 primary metabolites. Metabolites were identified based on the Level of Identification Standard of the Metabolomics Standards Initiative (Sumner et al., 2007).

### Transcriptome analysis

Using the same material as described in the metabolome analysis, total RNA was extracted from six cybrids, three in a Ler and three in an Ely nuclear background: L*er*^L*er*^ Ler^Ely^, *Ler^Bur^* and Ely^L*er*^ Ely^Ely^, Ely^Bur^ with three replicates per genotype, totaling 18 plants. Library preparation was done with a selection on 3’ polyadenylated tails to preferentially include nuclear mRNA. Read alignment was done using TopHat (Trapnell et al., 2009). Any chloroplast and mitochondrial genes remaining were excluded from further analysis. The raw counts were normalized and analyzed using the DeSeq2 package in R (Love et al., 2014). Genes for which the expression levels were significantly different between two cybrids were determined by comparing two genotypes using the contrast function of DeSeq2. P-values were determined using the Wald test, and p-values were adjusted using the Benjamini-Hochberg correction (α=0.05). GO enrichment analysis was done using default setting in g:profiler (g:GOSt). The complete set of detected genes in each cybrid was used as a statistical background in the analysis (Reimand et al., 2016).

### Phenotypic data analysis

We used the self-cybrids as our baseline in phenotypic comparisons to control for any possible effects of cybrid creation, with the exception of Bur^Bur^ which was replaced in all analysis with Bur-WT. Raw data was directly analyzed except for time series data of growth and chlorophyll reflectance which was preprocessed as follows. Time series data were fitted with a smooth spline using the gam function from the mgcv package in R (Wood et al., 2016). The fitted B-spline was subsequently used to derive curve parameters. These include area under the curve, slope under mean, first, second (median) and third quartile, minimal and maximal slope, and the timepoint where the slope is maximum. These parameters allow us to quantify not only plant size and growth rate but also the dynamic properties of the growth curve, i.e. did growth occur early of late, or was it more uniform? In addition, we calculated relative growth rate per time point by dividing the growth rate, relative to the plant size (Flood et al., 2016). All raw parameters and derived parameters were analyzed by fitting either a linear mixed model or a linear model. The linear mixed model was used when a random correction parameter was present, when such random correction parameters were absent a linear model was used. The models were analyzed using the Restricted Maximum Likelihood (REML) procedure for each relevant phenotype using the lme4 package in R (Bates et al., 2015). As each experiment had a different design, several models were employed (Supplementary Table 4). The following model was generally used, in some instances random terms (<u>underlined below</u>) were added:

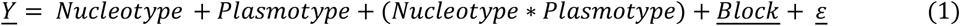

For every model, normality and equal variances were checked. Next for every phenotypic parameter we calculated significant difference, for the plasmotype and interaction term of the model (equation 1). This was done by ANOVA in which Kenward-Roger approximation for degrees of freedom was used. As posthoc tests we used a two-sided Dunnett’s test, where we tested whether a given cybrid was different from the self-cybrid control, within one nucleotype. Two side Hochberg’s posthoc tests were used when all pairwise comparisons were tested within one nucleotype (to test for epistasis) and across all nucleotypes (to test for additivity). The significance threshold for all posthoc tests was set at α=0.05. The contribution of the nucleotype, plasmotype and the interaction between the two, was determined by estimating the variance components in mixed models containing the same terms as in model (1). However, the fixed terms were taken as random:

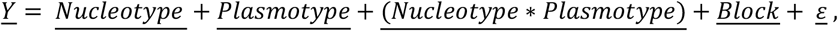

Where the variance components were estimated by the VarCorr function from the lme4 package. Total variance was calculated by summing all the variance components, after which the fraction explained variance for every term in the model was calculated. The broad sense heritability, in our case equal to repeatability (Falconer and Mackay, 1996), is determined by the three genetic components, i.e. nucleotype, plasmotype and their interaction. The fraction of broad sense heritability explained by the separate genetic components was calculated subsequently.

In total we measured 1859 phenotypes. After data processing, further analysis was only conducted on phenotypes with a broad sense heritability higher than 5%, removing phenotypes that were non-informative, leaving with 1782 phenotypes. Furthermore, to avoid biases in the results due to overly correlated data when stating summary statistics, we further subset the remaining 1782 phenotypes (Supplementary Data 2). Using a threshold based purely on correlation would favor the inclusion of variation largely driven by the nucleotype. Because the population is balanced, we therefore subtracted the averages of the nucleotype values from the cybrid phenotype values, to reveal the plasmotype effect per cybrid. From these we calculated the pearson correlations for all phenotypes. This highlighted that the most uncorrelated phenotypes mainly stem from one experiment assessing photosynthetic parameters under fluctuating light. The unbiased selection of a subset of phenotypes would result in the omission of several phenotypic categories. To present a balanced overview of all phenotypic categories we manually selected a subset comprising the following phenotypes. For time series in which we scored for up to 25 days after germination, we selected the morning measurements of day 8, 13, 18 and 23. The time series analysis of fluctuating light were measured for three (first experiment, Fig. 3) and five days (replicate experiment; Supplementary Fig. 8) in a row, with each day subjected to a different treatment. As these treatments reached their extremes in the middle and at end of the day, and the results of replicate experiments were very similar, we selected time points in the middle and at the end of the day of only the first experiment. For the different seed treatments we used the germination time until 50% of the seeds germinated. In addition, we included biomass, leaf movement, seed size, flowering time as single phenotypes and all 36 primary metabolites. This resulted in 92 phenotypes, that are used for giving summary- and test statistics (for a correlation plot of these, please see Supplementary Fig. 6). All data on the 1859 phenotypes, with summary- and test statistics are available in Supplementary Data 1 and Supplementary Table 3.

The correlation between plasmotype additive and plasmotype epistatic effects was calculated both with and without the Ely plasmotype. For both additive and epistatic effects every significant change between plasmotypes, within one nucleotype background, was counted (Supplementary Data 2). The Pearson correlation coefficients and accompanying p-values were calculated using the ggpubr package in R.

## Acknowledgements

:Hetty Blankestijn, Jose van de Belt, Daniel Oberste-Lehn, Elio Schijlen, Corrie Hanhart, and Joris ter Riele (all Wageningen University & Research) are acknowledged for help with experiments, Jonas Klasen (Max Planck Institute for Plant Breeding Research) for statistical advice, and Duur Aanen (Wageningen University & Research) for helpful discussions.

## Author contributions

P.J.F. and E.W. conceived and designed the study. T.P.J.M.T. designed and performed the statistical analysis with help from P.J.F., W.K. and F.v.E.. P.J.F., T.P.J.M.T., E.K., F. F.M.B., L.W., V.C.B., J.v.A., J.M.G., and L.S. performed experiments. P.J.F., T.P.J.M.T., K.S., P.K., E.S., J.A.H., S.K.S., R.W., W.L., R.M., F.v.E. and E.W. analysed data. D.M.K., J.J.B.K., M.K., J.H. and M.G.M.A. contributed to the interpretation of results. P.J.F., T.P.J.M.T. and E.W. wrote the paper with significant contributions from M.K., J.H. and M.G.M.A. All authors read and approved the final manuscript.

## Data availability

Sequencing and transcriptome data will be available in the European Nucleotide Archive with the primary accession code PRJEB29654. The raw datasets will be made available through Dryad, a reporting summary will be provided. The analysed datasets that support our findings are available as supplementary datasets. The associated raw data for Fig. 2 and 3 are provided in Supplementary Data 1, the raw data for Table 1 is provided in Supplementary Data 2. The germplasm generated in this project will be available via NASC.

## References

Bates D, Mächler M, Bolker B, Walker S (2015) Fitting linear mixed-effects models using lme4. Journal of Statistical Software 67: 48

Bock DG, Andrew RL, Rieseberg LH (2014) On the adaptive value of cytoplasmic genomes in plants. Mol. Ecol. 23: 4899–4911

Carreno-Quintero N, Acharjee A, Maliepaard C, Bachem CWB, Mumm R, Bouwmeester H, Visser RGF, Keurentjes JJB (2012) Untargeted Metabolic Quantitative Trait Loci Analyses Reveal a Relationship between Primary Metabolism and Potato Tuber Quality. Plant Physiology 158: 1306–1318

Chan KX, Phua SY, Crisp P, McQuinn R, Pogson BJ (2016) Learning the Languages of the Chloroplast: Retrograde Signaling and Beyond. 67: 25–53

Cingolani P, Platts A, Wang LL, Coon M, Nguyen T, Wang L, Land SJ, Lu X, Ruden DM (2012) A program for annotating and predicting the effects of single nucleotide polymorphisms, SnpEff. Fly 6: 80–92

Cingolani P, Platts A, Wang LL, Coon M, Nguyen T, Wang L, Land SJ, Lu X, Ruden DM (2012) A program for annotating and predicting the effects of single nucleotide polymorphisms, SnpEff: SNPs in the genome of Drosophila melanogaster strain w(1118); iso-2; iso-3. Fly 6: 80–92

Cruz JA, Savage LJ, Zegarac R, Hall CC, Satoh-Cruz M, Davis GA, Kovac WK, Chen J, Kramer DM (2016) Dynamic Environmental Photosynthetic Imaging Reveals Emergent Phenotypes. Cell Systems 2: 365–377

Dobler R, Rogell B, Budar F, Dowling DK (2014) A meta-analysis of the strength and nature of cytoplasmic genetic effects. J. Evolution Biol. 27: 2021–2034

Dowling DK, Abiega KC, Arnqvist G (2007) TEMPERATURE-SPECIFIC OUTCOMES OF CYTOPLASMIC-NUCLEAR INTERACTIONS ON EGG-TO-ADULT DEVELOPMENT TIME IN SEED BEETLES. 61: 194–201

El-Lithy ME, Rodrigues GC, van Rensen JJS, Snel JFH, Dassen HJHA, Koornneef M, Jansen MAK, Aarts MGM, Vreugdenhil D (2005) Altered photosynthetic performance of a natural *Arabidopsis* accession is associated with atrazine resistance. J. Exp. Bot. 56: 1625–1634

Falconer D, Mackay TJH, Essex, UK: Longmans Green (1996) Introduction to quantitative genetics. 1996. 3

Flood PJ, Harbinson J, Aarts MGM (2011) Natural genetic variation in plant photosynthesis. Trends Plant Sci. 16: 327–335

Flood PJ, Kruijer W, Schnabel SK, Schoor R, Jalink H, Snel JFH, Harbinson J, Aarts MGM (2016) Phenomics for photosynthesis, growth and reflectance in Arabidopsis thaliana reveals circadian and long-term fluctuations in heritability. Plant Methods 12: 1–14

Flood Pádraic J, van Heerwaarden J, Becker F, de Snoo CB, Harbinson J, Aarts Mark GM (2016) Whole-Genome Hitchhiking on an Organelle Mutation. Current Biology 26: 1306–1311

Flood PJ, Yin L, Herdean A, Harbinson J, Aarts MGM, Spetea C (2014) Natural variation in phosphorylation of photosystem II proteins in Arabidopsis thaliana: is it caused by genetic variation in the STN kinases? Philosophical Transactions of the Royal Society B: Biological Sciences 369

Gobron N, Waszczak C, Simon M, Hiard S, Boivin S, Charif D, Ducamp A, Wenes E, Budar F (2013) A Cryptic Cytoplasmic Male Sterility Unveils a Possible Gynodioecious Past for *Arabidopsis thaliana*. PLoS ONE 8: e62450

Hill GE, Havird JC, Sloan DB, Burton RS, Greening C, Dowling DK (2019) Assessing the fitness consequences of mitonuclear interactions in natural populations. 94: 1089–1104

Hoekstra LA, Siddiq MA, Montooth KL (2013) Pleiotropic Effects of a Mitochondrial–Nuclear Incompatibility Depend upon the Accelerating Effect of Temperature in Drosophila. 195: 1129–1139

Houben A, Sanei M, Pickering R (2011) Barley doubled-haploid production by uniparental chromosome elimination. 104: 321–327

Jin S, Daniell H (2015) The Engineered Chloroplast Genome Just Got Smarter. Trends in Plant Science 20: 622–640

Joosen RVL, Kodde J, Willems LAJ, Ligterink W, van der Plas LHW, Hilhorst HWM (2010) germinator: a software package for high-throughput scoring and curve fitting of Arabidopsis seed germination. The Plant Journal 62: 148–159

Joseph B, Corwin JA, Li B, Atwell S, Kliebenstein DJ (2013) Cytoplasmic genetic variation and extensive cytonuclear interactions influence natural variation in the metabolome. eLife 2: e00776

Joseph B, Corwin JA, Li B, Atwell S, Kliebenstein DJ (2013) Cytoplasmic genetic variation and extensive cytonuclear interactions influence natural variation in the metabolome, Vol 2

Joseph B, Corwin JA, Züst T, Li B, Iravani M, Schaepman-Strub G, Turnbull LA, Kliebenstein DJ (2013) Hierarchical Nuclear and Cytoplasmic Genetic Architectures for Plant Growth and Defense within Arabidopsis. The Plant Cell Online 25: 1929–1945

Karimi-Ashtiyani R, Ishii T, Niessen M, Stein N, Heckmann S, Gurushidze M, Banaei-Moghaddam AM, Fuchs J, Schubert V, Koch K, Weiss O, Demidov D, Schmidt K, Kumlehn J, Houben A (2015) Point mutation impairs centromeric CENH3 loading and induces haploid plants. 112: 11211–11216

Kermicle JL (1969) Androgenesis Conditioned by a Mutation in Maize. 166: 1422–1424

Kleine T, Leister D (2016) Retrograde signaling: Organelles go networking. Biochimica et Biophysica Acta (BBA) - Bioenergetics 1857: 1313–1325

Kokorian J, Polder G, Keurentjes JJB, Vreugdenhil D, Olortegui Guzman MC (2010) An ImageJ based measurement setup for automated phenotyping of plants. In A Jahnen, C Moll, eds, Proceedings of the ImageJ User and Developer Conference, Luxembourg, Luxembourg, 27-29 October 2010. Centre de Recherche Public Henri Tudor, Luxembourg, pp 178–182

Kromdijk J, Głowacka K, Leonelli L, Gabilly ST, Iwai M, Niyogi KK, Long SP (2016) Improving photosynthesis and crop productivity by accelerating recovery from photoprotection. Science 354: 857–861

Kwak S-Y, Lew TTS, Sweeney CJ, Koman VB, Wong MH, Bohmert-Tatarev K, Snell KD, Seo JS, Chua N-H, Strano MS (2019) Chloroplast-selective gene delivery and expression in planta using chitosan-complexed single-walled carbon nanotube carriers. Nature Nanotechnology 14: 447–455

Levings CS (1990) The Texas Cytoplasm of Maize: Cytoplasmic Male Sterility and Disease Susceptibility. Science 250: 942–947

Li H (2013) Aligning sequence reads, clone sequences and assembly contigs with BWA-MEM. In arXiv e-prints,

Li H, Handsaker B, Wysoker A, Fennell T, Ruan J, Homer N, Marth G, Abecasis G, Durbin R, Subgroup GPDP (2009) The Sequence Alignment/Map format and SAMtools. Bioinformatics 25: 2078–2079

Lisec J, Schauer N, Kopka J, Willmitzer L, Fernie AR (2006) Gas chromatography mass spectrometry-based metabolite profiling in plants. Nat. Protocols 1: 387–396

Love MI, Huber W, Anders S (2014) Moderated estimation of fold change and dispersion for RNA-seq data with DESeq2. Genome Biology 15: 550

Martin M (2011) Cutadapt removes adapter sequences from high-throughput sequencing reads. 2011 17: 3 %J EMBnet.journal

Miclaus M, Balacescu O, Has I, Balacescu L, Has V, Suteu D, Neuenschwander S, Keller I, Bruggmann R (2016) Maize Cytolines Unmask Key Nuclear Genes That Are under the Control of Retrograde Signaling Pathways in Plants. Genome Biology and Evolution 8: 3256–3270

Montooth KL, Meiklejohn CD, Abt DN, Rand DM (2010) MITOCHONDRIAL-NUCLEAR EPISTASIS AFFECTS FITNESS WITHIN SPECIES BUT DOES NOT CONTRIBUTE TO FIXED INCOMPATIBILITIES BETWEEN SPECIES OF DROSOPHILA. Evolution 64: 3364–3379

Mossman JA, Biancani LM, Zhu C-T, Rand DM (2016) Mitonuclear Epistasis for Development Time and Its Modification by Diet in <em>Drosophila</em>. 203: 463–484

Mossman JA, Ge JY, Navarro F, Rand DM (2019) Mitochondrial DNA Fitness Depends on Nuclear Genetic Background in <em>Drosophila</em>. 9: 1175–1188

Murchie EH, Kefauver S, Araus JL, Muller O, Rascher U, Flood PJ, Lawson T (2018) Measuring the dynamic photosynthome. Annals of Botany 122: 207–220

Peterson R, Slovin JP, Chen CJIJoPB (2010) A simplified method for differential staining of aborted and non-aborted pollen grains. 1: e13–e13

Petrillo E, Godoy Herz MA, Fuchs A, Reifer D, Fuller J, Yanovsky MJ, Simpson C, Brown JWS, Barta A, Kalyna M, Kornblihtt AR (2014) A Chloroplast Retrograde Signal Regulates Nuclear Alternative Splicing. Science 344: 427–430

Ravi M, Chan SWL (2010) Haploid plants produced by centromere-mediated genome elimination. 464: 615–618

Ravi M, Marimuthu MPA, Tan EH, Maheshwari S, Henry IM, Marin-Rodriguez B, Urtecho G, Tan J, Thornhill K, Zhu F, Panoli A, Sundaresan V, Britt AB, Comai L, Chan SWL (2014) A haploid genetics toolbox for Arabidopsis thaliana. Nat Commun 5

Reimand J, Arak T, Adler P, Kolberg L, Reisberg S, Peterson H, Vilo J (2016) g:Profiler—a web server for functional interpretation of gene lists (2016 update). Nucleic Acids Research 44: W83–W89

Roux F, Mary-Huard T, Barillot E, Wenes E, Botran L, Durand S, Villoutreix R, Martin-Magniette M-L, Camilleri C, Budar F (2016) Cytonuclear interactions affect adaptive traits of the annual plant *Arabidopsis thaliana* in the field. Proceedings of the National Academy of Sciences 113: 3687–3692

Ruf S, Forner J, Hasse C, Kroop X, Seeger S, Schollbach L, Schadach A, Bock R (2019) High-efficiency generation of fertile transplastomic Arabidopsis plants. Nature Plants 5: 282–289

Sambatti JBM, Ortiz-Barrientos D, Baack EJ, Rieseberg LH (2008) Ecological selection maintains cytonuclear incompatibilities in hybridizing sunflowers. 11: 1082–1091

Schneerman M, Charbonneau M, Weber DJMGCN (2000) A survey of ig containing materials. 54–55

Sloan DB, Wu Z, Sharbrough J (2018) Correction of Persistent Errors in Arabidopsis Reference Mitochondrial Genomes. 30: 525–527

Somerville CR, Ogren WL (1980) Photorespiration mutants of *Arabidopsis thaliana* deficient in serine-glyoxylate aminotransferase activity. Proceedings of the National Academy of Sciences 77: 2684–2687

Sumner LW, Amberg A, Barrett D, Beale MH, Beger R, Daykin CA, Fan TW-M, Fiehn O, Goodacre R, Griffin JL, Hankemeier T, Hardy N, Harnly J, Higashi R, Kopka J, Lane AN, Lindon JC, Marriott P, Nicholls AW, Reily MD, Thaden JJ, Viant MR (2007) Proposed minimum reporting standards for chemical analysis. Metabolomics 3: 211–221

Tang Z, Hu W, Huang J, Lu X, Yang Z, Lei S, Zhang Y, Xu C (2014) Potential Involvement of Maternal Cytoplasm in the Regulation of Flowering Time via Interaction with Nuclear Genes in Maize. Crop Sci. 54: 544–553

The 1001 Genomes Consortium (2016) 1,135 Genomes Reveal the Global Pattern of Polymorphism in Arabidopsis thaliana. Cell

Trapnell C, Pachter L, Salzberg SL (2009) TopHat: discovering splice junctions with RNA-Seq. Bioinformatics 25: 1105–1111

Wehrens R, Hageman JA, van Eeuwijk F, Kooke R, Flood PJ, Wijnker E, Keurentjes JJB, Lommen A, van Eekelen HDLM, Hall RD, Mumm R, de Vos RCH (2016) Improved batch correction in untargeted MS-based metabolomics. Metabolomics 12: 88

Wijnker E, Deurhof L, van de Belt J, de Snoo CB, Blankestijn H, Becker F, Ravi M, Chan SWL, van Dun K, Lelivelt CLC, de Jong H, Dirks R, Keurentjes JJB (2014) Hybrid recreation by reverse breeding in Arabidopsis thaliana. Nature protocols 9: 761–772

Wood SN, Pya N, Säfken B (2016) Smoothing Parameter and Model Selection for General Smooth Models. Journal of the American Statistical Association 111: 1548–1563

Yin L, Fristedt R, Herdean A, Solymosi K, Bertrand M, Andersson MX, Mamedov F, Vener AV, Schoefs B, Spetea C (2012) Photosystem II Function and Dynamics in Three Widely Used *Arabidopsis thaliana* Accessions. PLoS ONE 7: e46206

Zeyl C, Andreson B, Weninck E (2005) NUCLEAR-MITOCHONDRIAL EPISTASIS FOR FITNESS IN SACCHAROMYCES CEREVISIAE. Evolution 59: 910–914

Zhang J, Khan SA, Hasse C, Ruf S, Heckel DG, Bock R (2015) Full crop protection from an insect pest by expression of long double-stranded RNAs in plastids. Science 347: 991–994

