## Supplemental Figures and Tables for "Reciprocal cybrids reveal how organellar genomes affect plant phenotypes"

**Supplementary Information:**

**Supplementary Data 1.** Summary and test statistics for all 1859 phenotypes. Tab 1 shows the least squared means for all phenotypes per cybrid, these are calculated via the models as specified in Supplementary Table 4. Tab 2 shows the fraction explained variance for the nucleotype, plasmotype and the interaction separately including cybrids with the Ely plasmotype. We show the variance components for all terms in the model. Using the variance components of the genetic components we calculated the broad sense heritability (H^2^). We used a H^2^ threshold of 0.05 for phenotypes to be included in summary- and test statistics. The fraction of broad sense heritability, for all three genetic components, was given as well. Tab 3 shows the same data as tab 2, but excluding the cybrids with the Ely plasmotype. Tab 4 shows the significant plasmotype additive effects, with Hochberg’s p-value correction. Letters indicate significance or not. Tab 5 shows the significant plasmotype epistatic effects. This was done for every plasmotype within each nucleotype (with Hochberg’s p-value correction), as well as the comparison between the self-cybrids and the other cybrids within each nucleotype (with Dunnett’s p-value correction).

**Supplementary Data 2.** Summary and test statistics of 92 phenotypes, with easy to use tables to search for significant differences for all separate phenotypes. Tab 1 shows the significant additive effects for every pairwise comparison. With a text box to explain the interpretation of the data presented. Tab 1 contains the underlying data used for generating Table 1A. Tab 2 gives the significant difference for every pairwise comparison within a nucleotype i.e. epistatic effects. A text box is provided to explain the interpretation of the data. Tab 2 contains the underlying data used for generating Table 1B. The remaining tabs show summary and test statistics for the 92 phenotypes, in the same manner as in Supplementary Data 1.

**Supplementary Data 3.** Differential expression overview from the RNAseq experiment for the 6 cybrid comparisons. Tabs 1 till 6 show pairwise comparisons of cybrid lines, indicated as “Nucleotype-Plasmotype vs Nucleotype-Plasmotype”. For every detected nuclear gene the differential expression data is given, with the adjusted p-value cut-off set at α=0.05 (in yellow). The summary statistics for this are given in Supplementary Table 5. Tab 7 shows the GO enrichment for 5 cybrid comparisons. For one comparison (L*er*^Bur^ vs L*er*^L^*^er^*) we only detected three significantly expressed genes for which a GO enrichment yielded no results. This tab also shows the GO enrichment for genes that differentially change expression when The Ely plasmotype is replaced by L*er* or Bur in L*er* and Ely nuclear backgrounds (indicated as “Ely main”).

**Supplementary Data 4.** Predicted impact of SNPs and INDELs of the chloroplastic and mitochondrial genomes of all 7 accessions used. For every variant the reference allele, based on TAIR10.1, is given next to the alternative allele. We indicated for all 7 accessions whether they share the reference (0/0) or alternative allele (1/1), and used SnpEff to predict the impact. The changes are ranked as “Low”, “Modifier”, “Moderate” and “High”, based on the location in respect to a gene, and predicted amino acid change. In our interpretation we have used the “Moderate” and “High” impact variants. The gene name affected, as well as the nucleotide and amino acid change are provided.

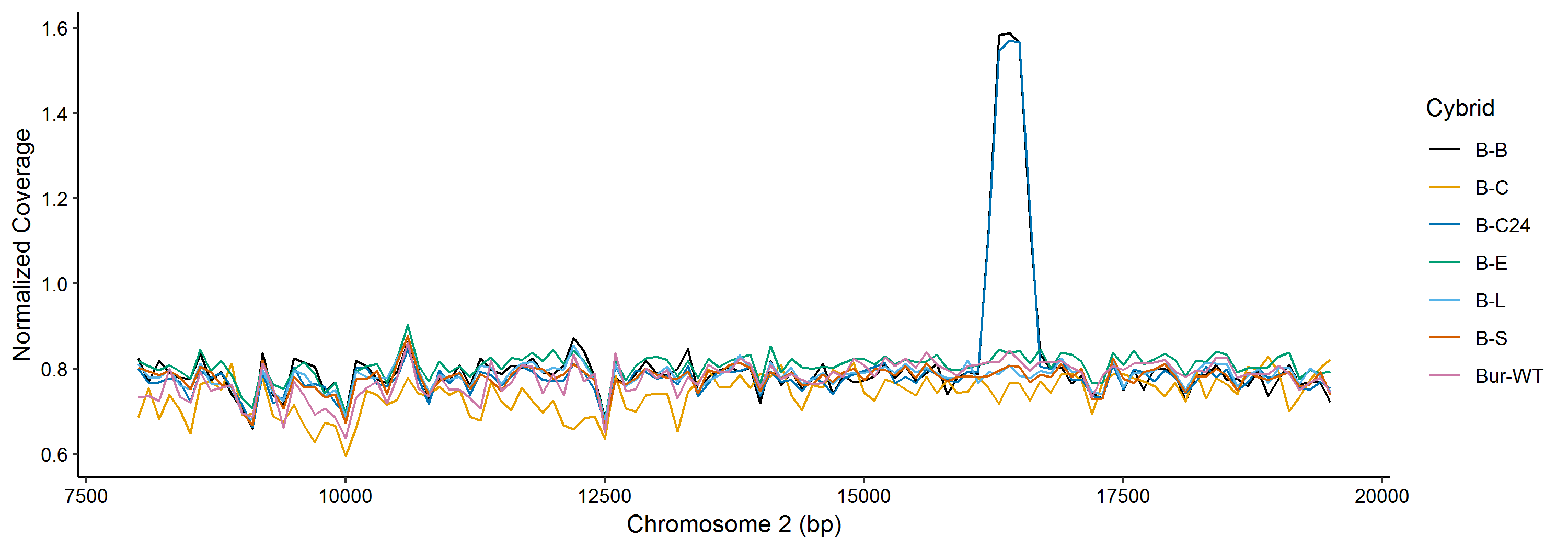
**Supplementary Figure 1.** Coverage plots for the long arm of chromosome 2 revealed the presence of a spontaneous nuclear DNA duplication in two cybrid lines (Bur^Bur^ and Bur^C24^). These lines were excluded from all further analyses.

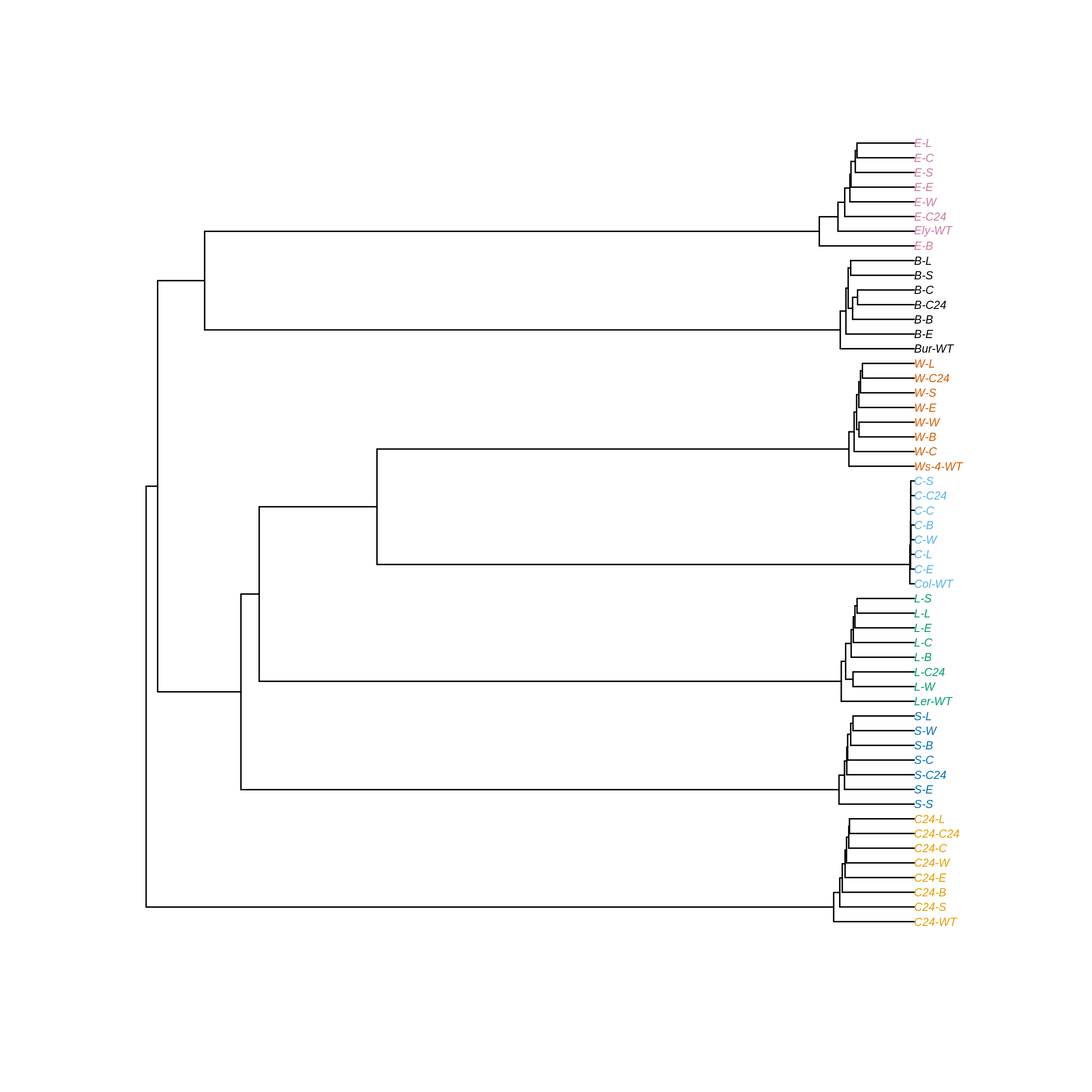

**Supplementary Figure 2**. Hierarchical clustering of the cybrids and wildtypes based on 3699159 nuclear SNPs spread equally over all chromosomes. The variation identified within a cluster of the same nucleotype is due to erroneous base calls induced by Illumina short read sequencing (on average below 0.0001% mismatches). Names at the tip of the branches indicate a nucleotype-plasmotype cybrid combination (Bur = B, C24 = C24, Col = C, L*er* = L, Sha = S, Ws-4 = W, Ely = E, Wild-type = WT). Samples are coloured according to nucleotype.

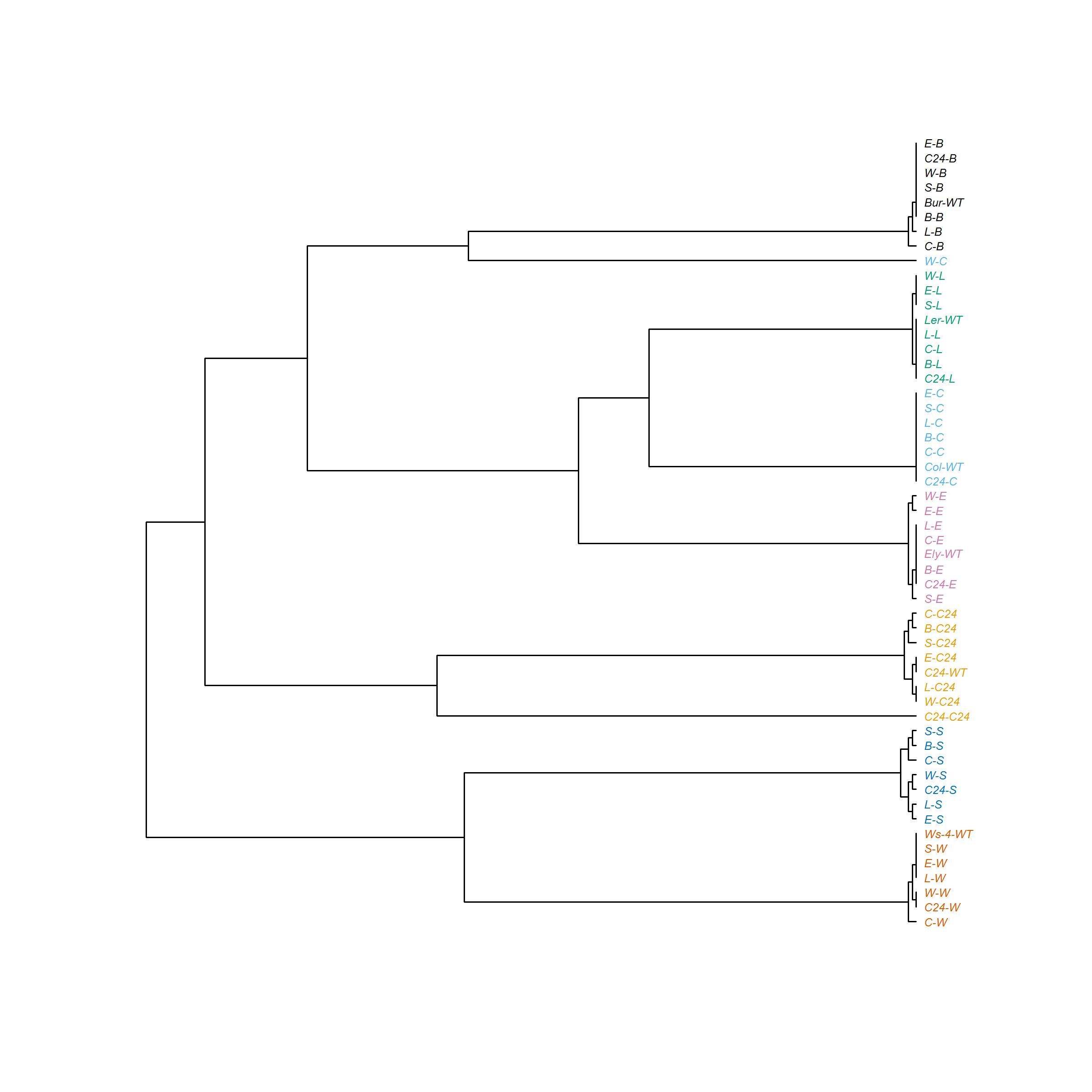
**Supplementary Figure 3.** Hierarchical clustering of the cybrids and wildtypes based on 356 chloroplast SNPs spread across the chloroplast genome. Names at the tip of the branches indicate specific nucleotype-plasmotype cybrid combinations. (Bur = B, C24 = C24, Col = C, L*er* = L, Sha = S, Ws-4 = W, Ely = E, Wild-type = WT). Sample names are coloured according to their plasmotypes. Sample W-C (i.e. WS-4^Col^) forms a cluster with cybrids containing the Bur plasmotype and, like C24-C24 (C24^C24^), has a very long branch. This is an artefact caused by sample contamination during sequencing. Please see online methods for further information.

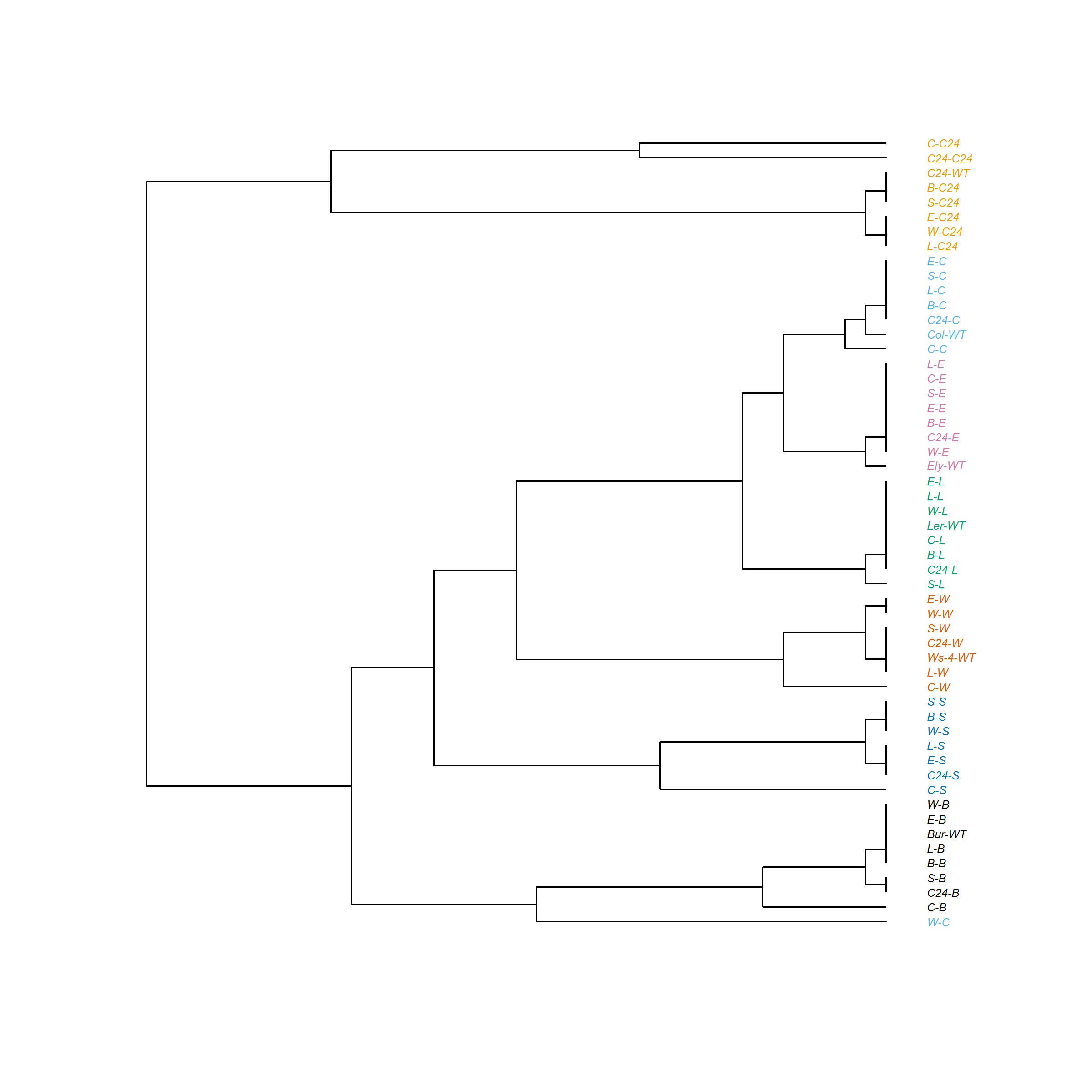
**Supplementary Figure 4.** Hierarchical clustering of the cybrids and wild types based on 135 mitochondrial SNPs spread across the mitochondrial genome. Names at the tip of the branches indicate a nucleotype-plasmotype cybrid combination (Bur = B, C24 = C24, Col = C, L*er* = L, Sha = S, Ws-4 = W, Ely = E, Wild-type = WT). Sample names are coloured according to their plasmotype. Sample W-C (i.e. WS-4^Col^) forms a cluster with cybrids containing the Bur plasmotype. This is an artefact caused by sample contamination during sequencing. Please see online methods for further information.

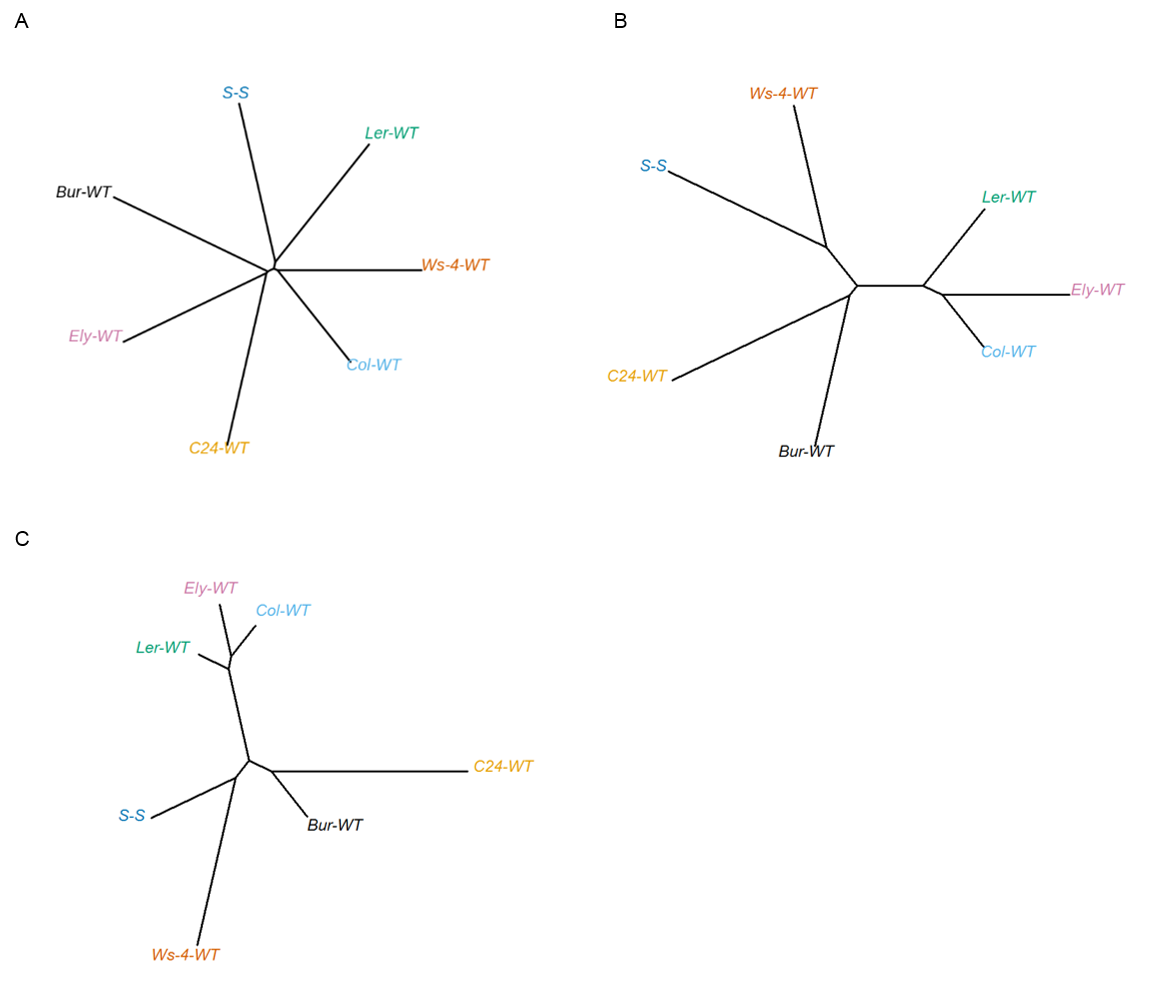
**Supplementary Figure 5.** Neighbor joining (NJ) trees based on nuclear and organellar genomes for the seven *Arabidopsis thaliana* accessions. A) NJ trees based on nuclear SNPs and INDELs B) NJ trees based on chloroplast SNPs and INDELs C) NJ trees based on mitochondrial SNPs and INDELs.

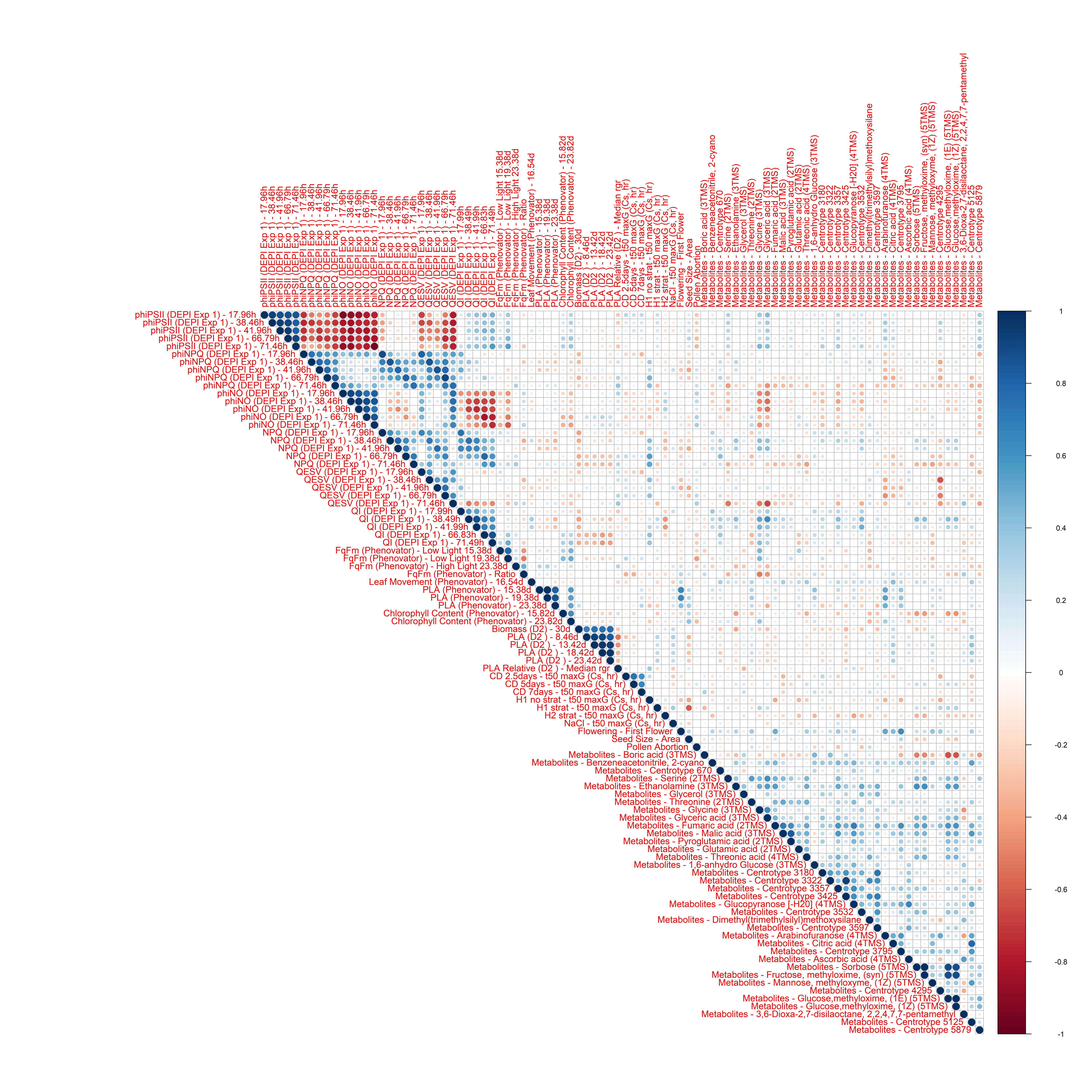

**Supplementary Figure 6.** Correlation plot based on the plasmotype effects of the subset of 92 phenotypes. For more information on the 92 phenotypes see Supplementary Table 2 and 4. Dark blue has a correlation of 1 and dark red a correlation of -1. Correlation represents the Pearson correlation coefficient.

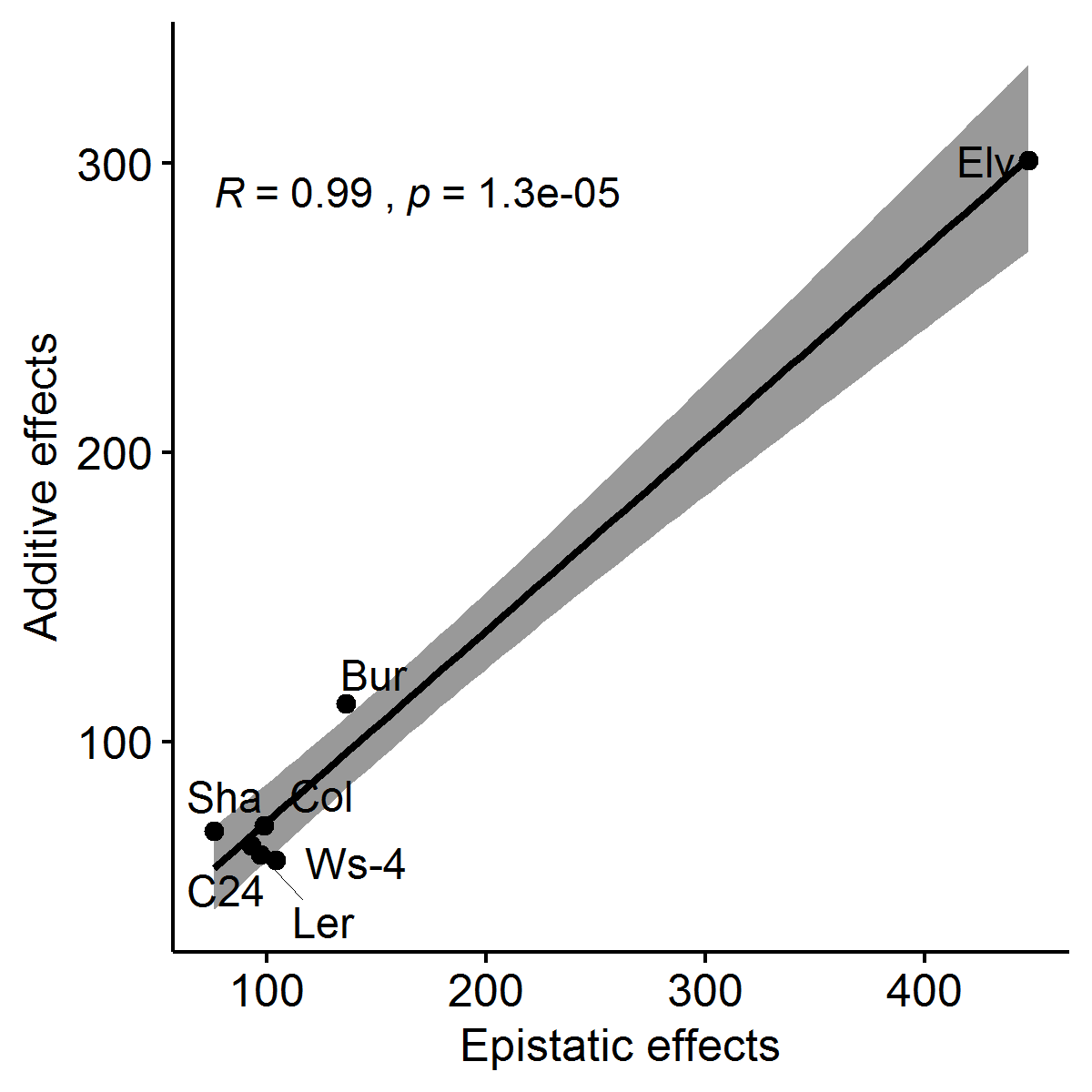

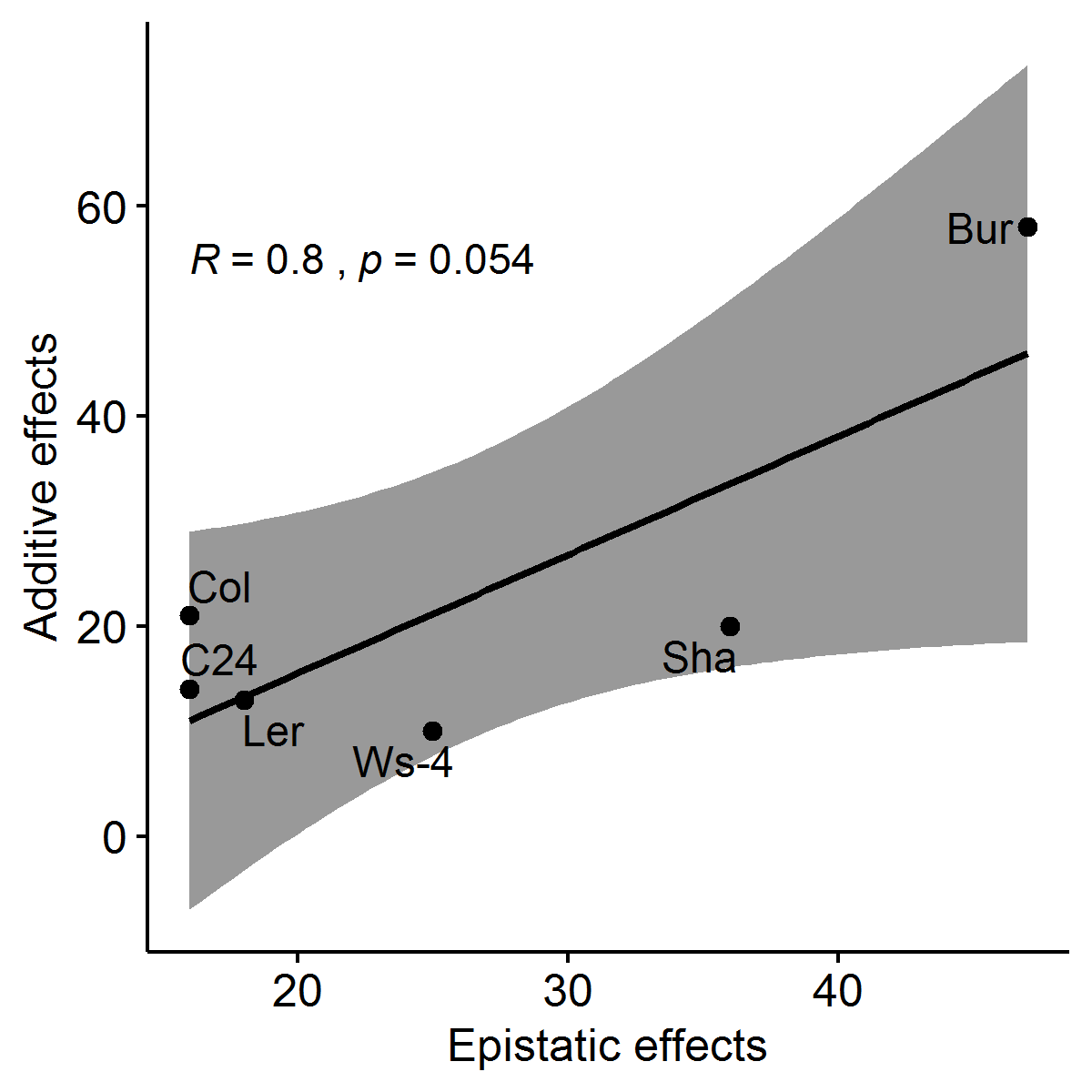

A

B

**Supplementary Figure 7.** Scatterplots showing the correlation between the number of plasmotype additive and plasmotype epistatic effects. A) showing the spread when including the Ely plasmotype in every comparison. B) showing the spread when excluding the Ely plasmotype. R represents the Pearson correlation coefficient.

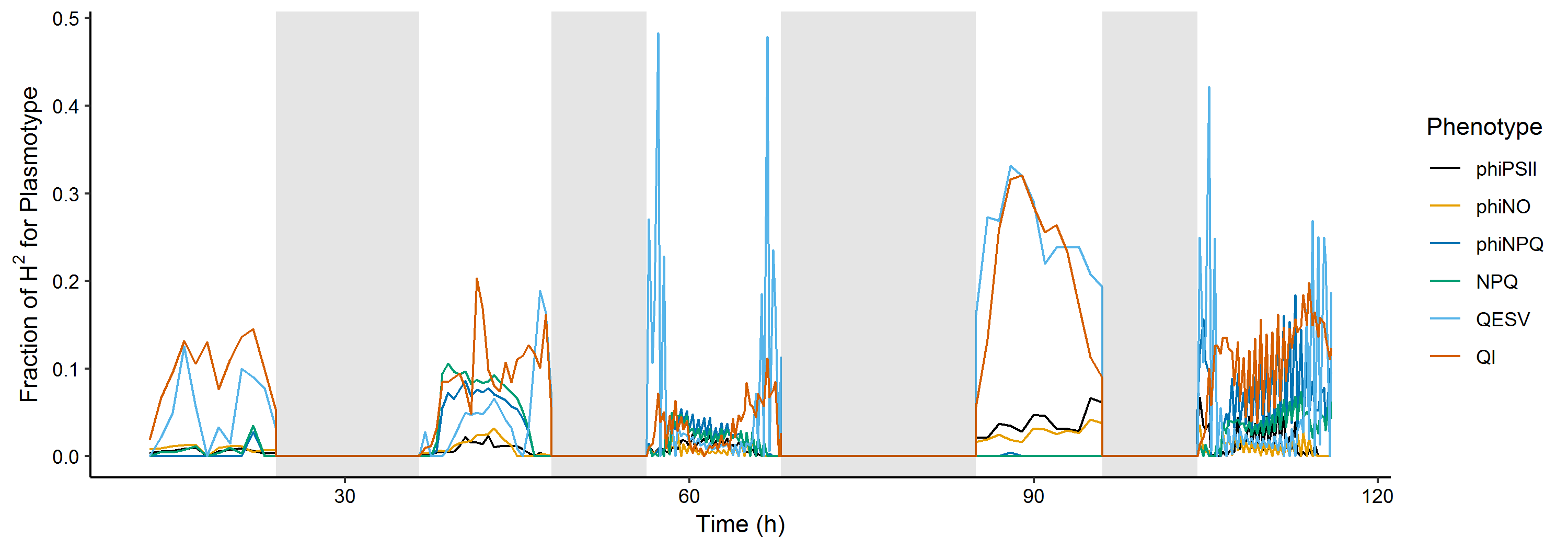

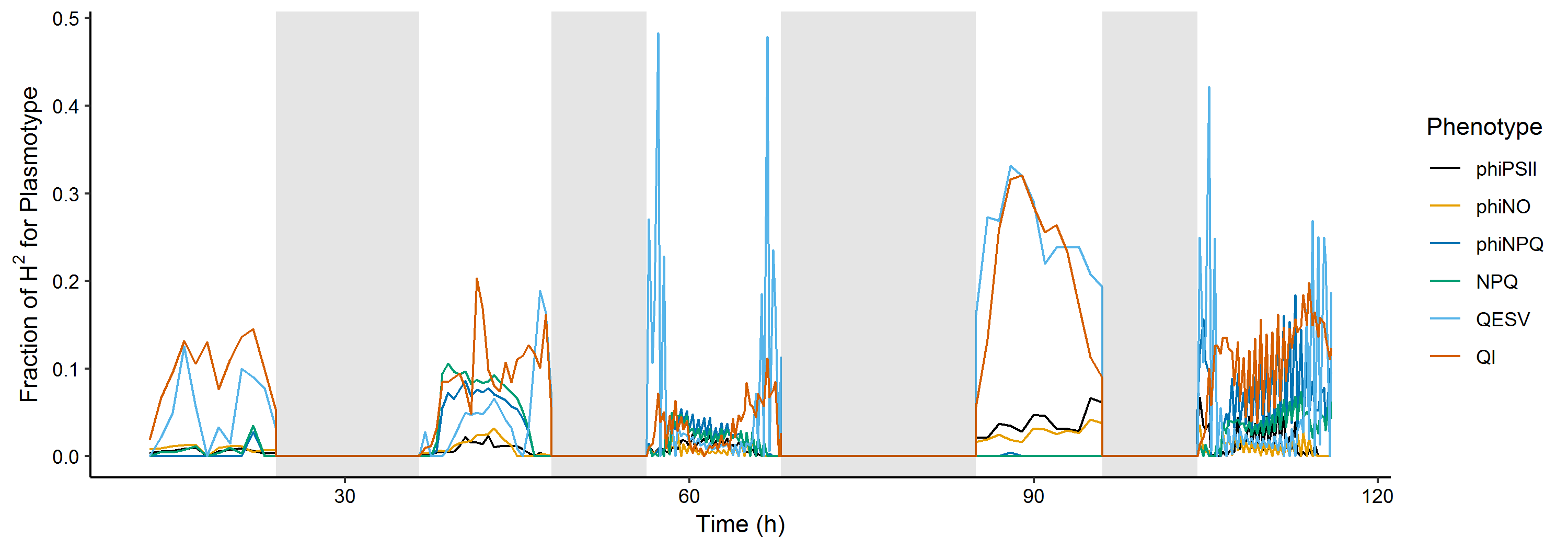

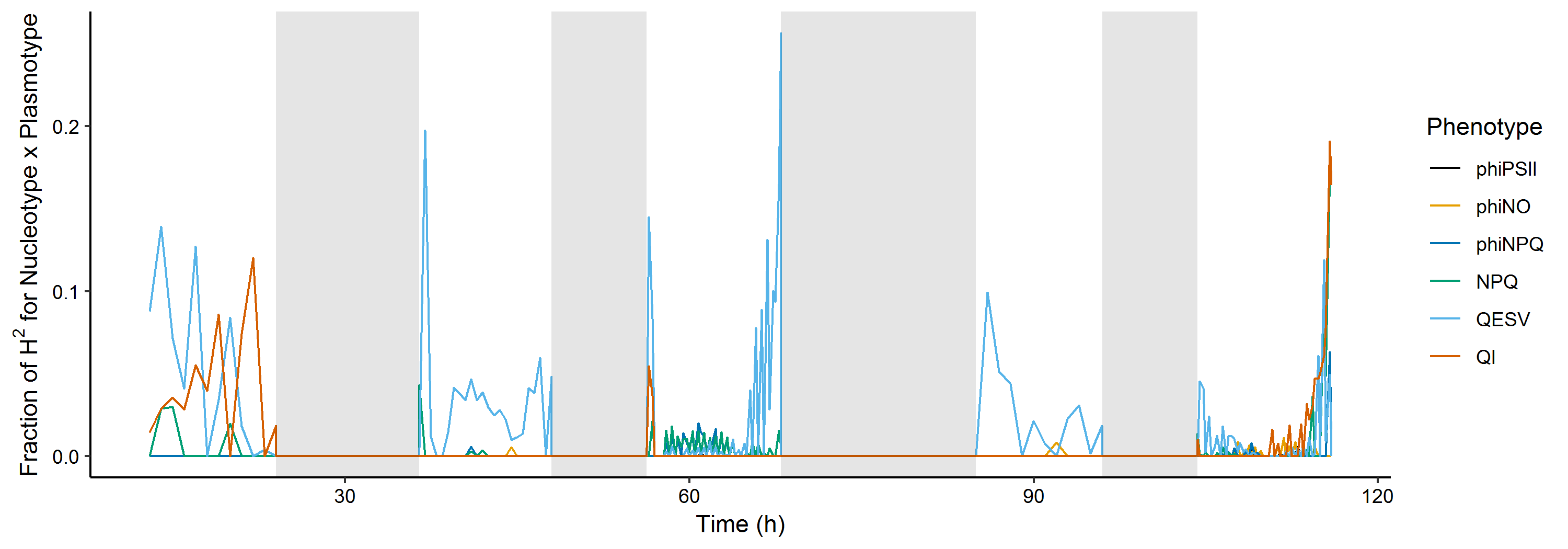

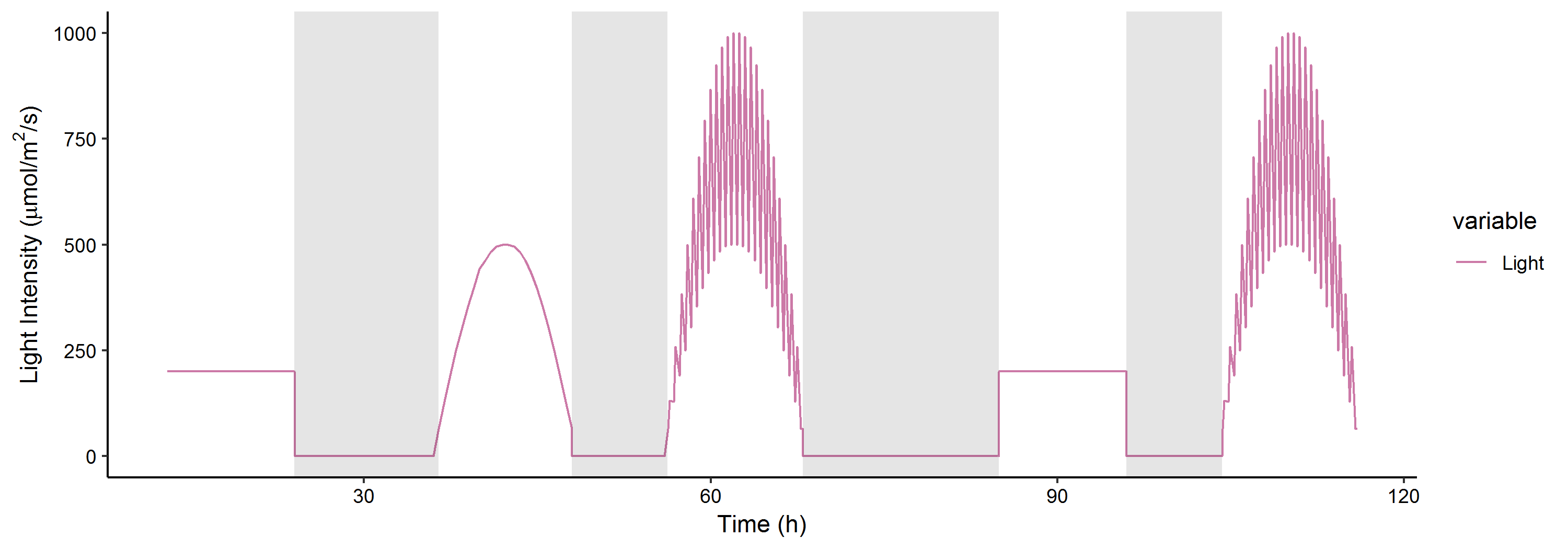

Φ_PSII_

Φ_NPQ_

Φ_NO_

NPQ

q_E_

q_I_

Phenotype

C

B

A

**Supplementary Figure 8.** The fraction of explained genetic variation (H^2^) for changes in photosynthesis phenotypes in response to light conditions. A) shows the fraction of H^2^ for epistatic interactions. B) panel shows H^2^ for plasmotype additive effects. C) shows the light intensity for five consecutive days with growth under: steady light (day 1); in- and decreasing light intensity (day 2); fluctuating in- and decreasing light intensity (day 3); steady light (day 4) and fluctuating in- and decreasing light intensity (day 5). Days are separated by nights (shaded areas). The first three days of this experiment are identical to the light conditions of the experiment shown in main Fig. 3.

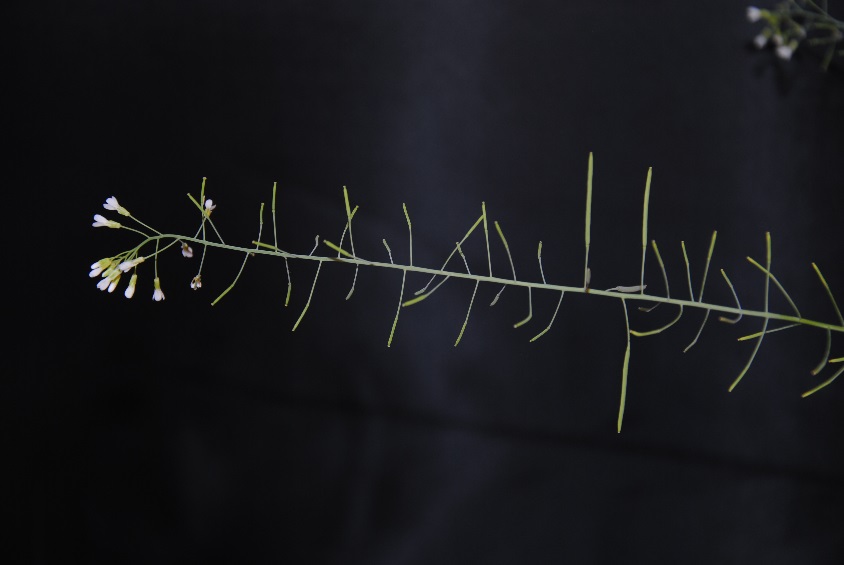

A

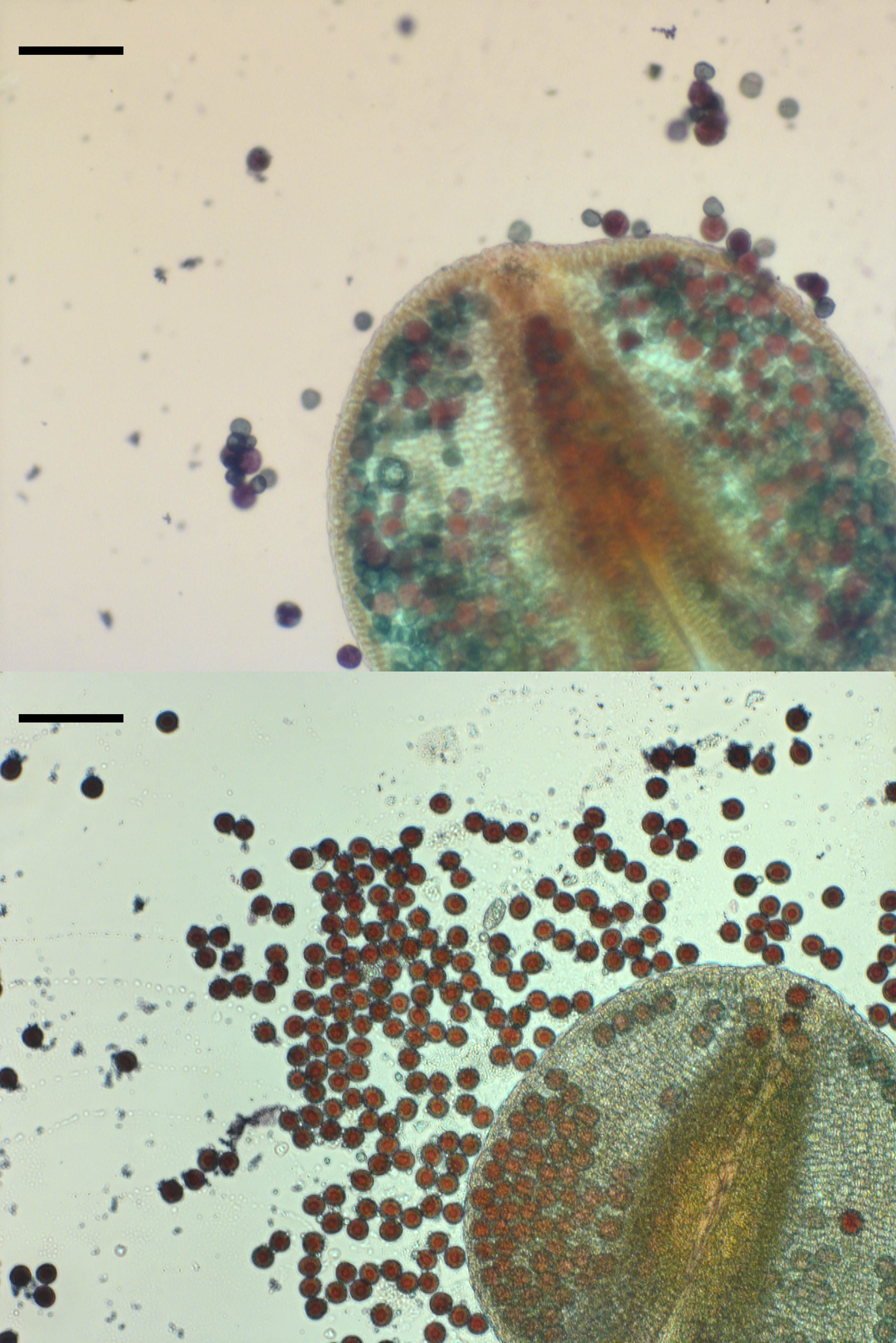

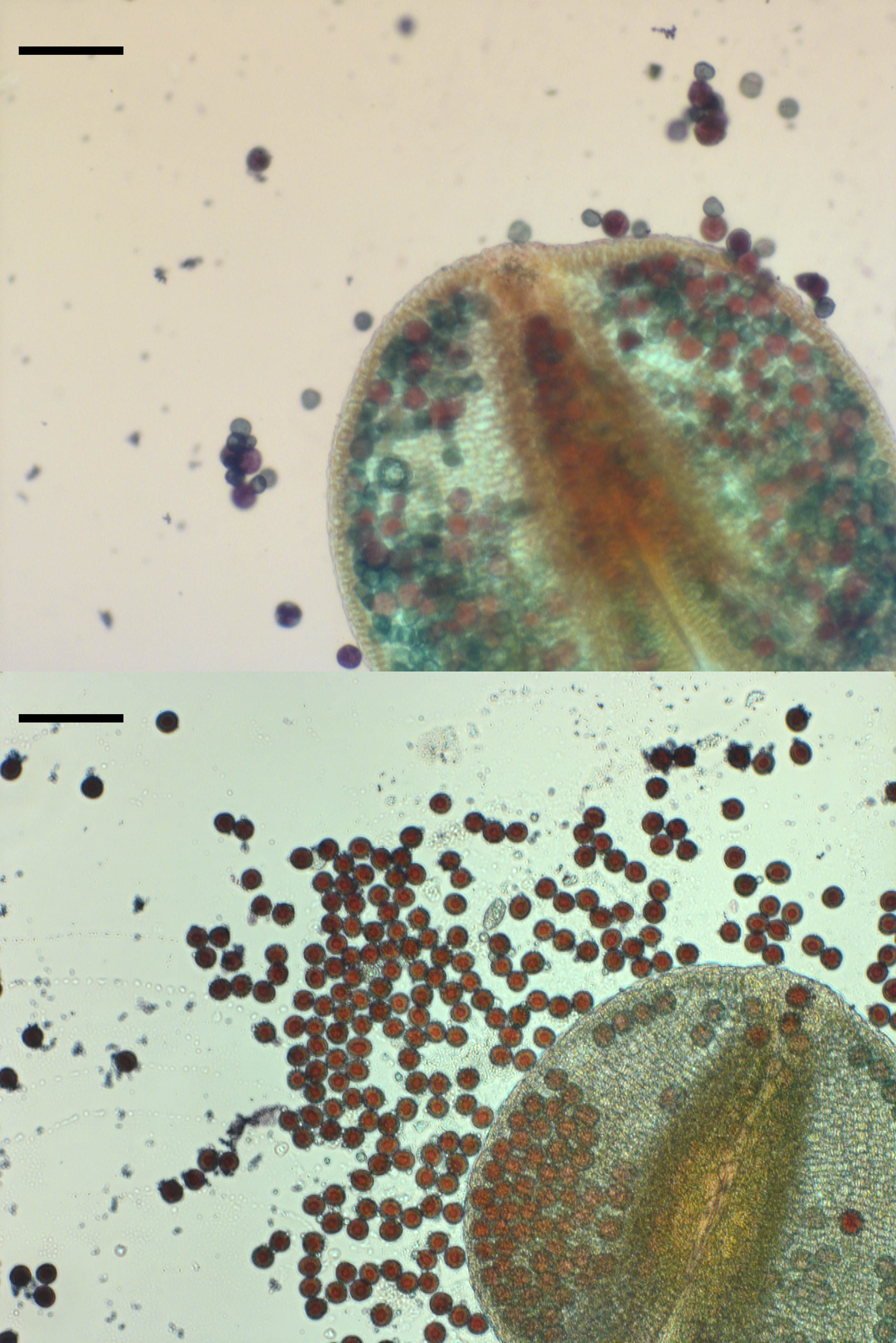

B

C

**Supplementary Figure 9.** Cytoplasmic male sterility in Ely^Sha^. A) An Ely^Sha^ plant was polinated on three open flowers using Ely wild-type pollen, which produced elongated siliques (indicated with red arrows). Scale bar is 1 cm. B) Anther and pollen of Ely^Sha^, stained with Alexander stain. Note the presence of a high percentage (45%; n=750) of greenish, almost colourless aborted pollen. Pollen with a red colour in this line are not able to fertilize ovules, as deduced from the male sterile phenotype of Ely^Sha^ (as shown in panel A). Scale bar is 500 μm. C) Anther and pollen of Ely wildtype. Note that all pollen have a dark red colour, suggesting high viability. Ely wildtype is able to fertilize Ely^Sha^ (as shown in panel A).

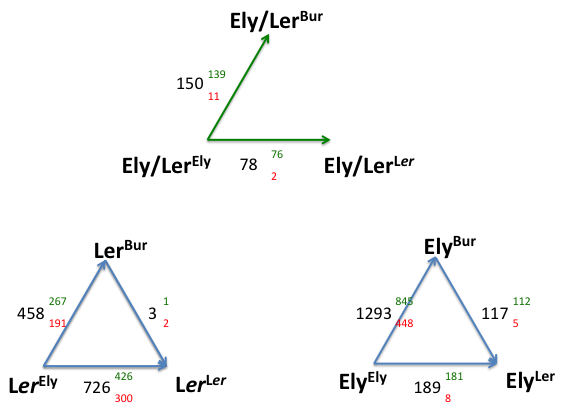

**Supplementary Figure 10.** Changes in gene expression between cybrids with a L*er* nucleus (lower left triangle) and an Ely nucleus (lower right triangle). Cybrid genotypes are indicated as Nucleotype^Plasmotype^. The triangle on the lower left shows cybrid comparisons with a L*er* nuclear background (for plasmotypes Ely, L*er* and Bur and the triangle on the lower right shows cybrid comparisons with an Ely nuclear background for the same plasmotypes. Significantly Differentially Expressed (DE) genes between cybrid comparisons are indicated with black numbers. These DE genes are subdivided in upregulated genes (green numbers in superscript) and downregulated genes (red numbers in subscript), following the direction of the arrows between cybrids (i.e. the change from an Ely to a L*er* plasmotype in a L*er* nuclear background resulted in 726 differentially expressed genes, of which 426 were upregulated and 300 were downregulated). The green upper triangle shows what differentially expressed genes the comparisons in the lower triangles have in common. For example, the L*er* and Ely nuclear backgrounds show of common responses of 78 DE genes when the Ely plasmotype is changed for a L*er* plasmotype. The absence of one of the comparisons in this triangle is due to the absence of shared DE genes. The common effect of changing an Ely plasmotype for either Bur of L*er* was derived by assessing what DE genes are similar along the axes in the green triangle. These 78 and 150 genes have 40 shared DE genes.

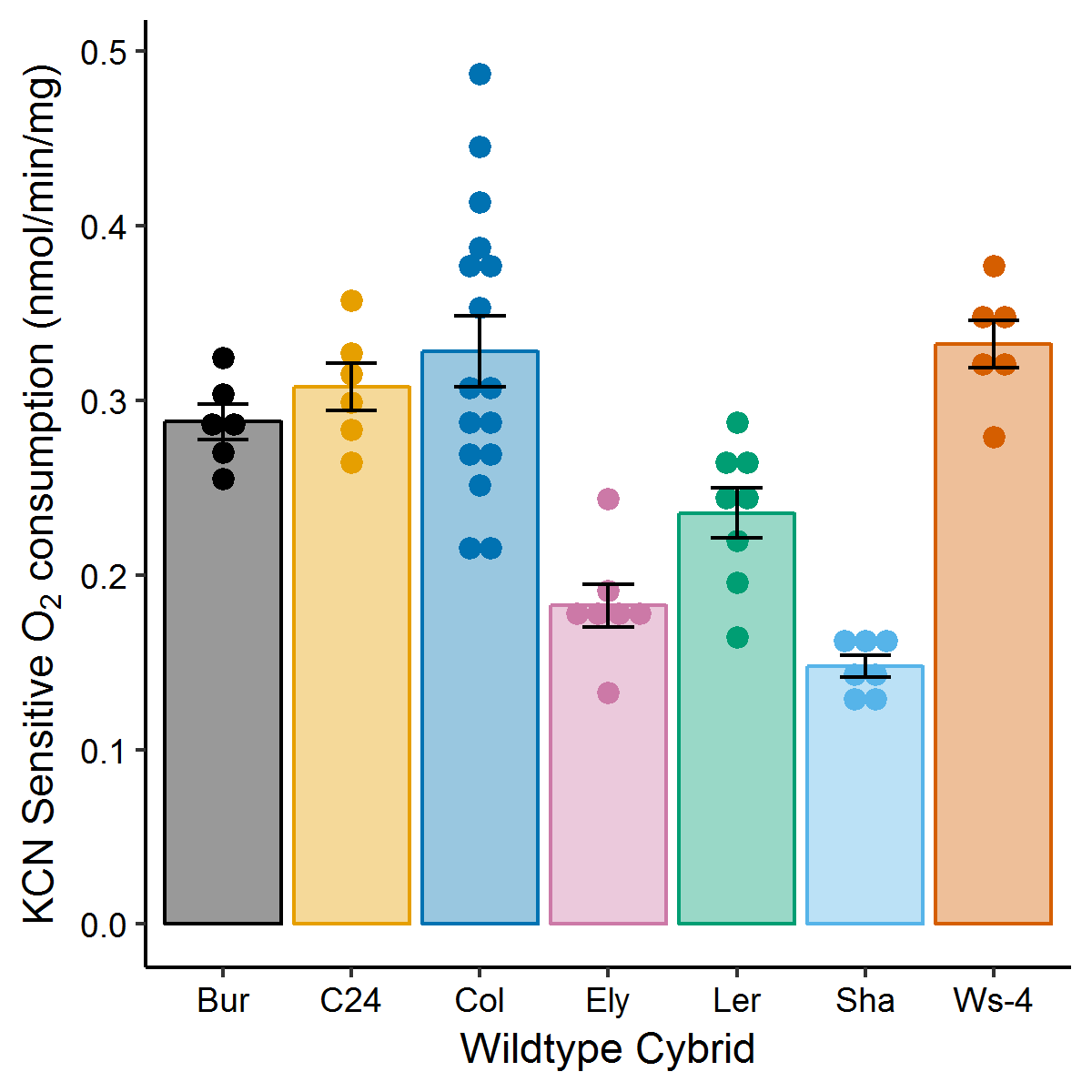

cd	cd	d	ab	bc	a	d

**Supplementary Figure 11.** KCN sensitive O_2_ consumption of mitochondrial seedlings of wild-type accessions. Mitochondrial ATP-synthesis proceeds mainly through the phosphorylating cytochrome (KCN sensitive) pathway. KCN sensitive O_2_ consumption by Bur does not differ significantly from C24, Col and Ws-4. Error bars represent the standard error of the mean (n=6-11). Letters indicate significant differences using the Tukey posthoc test.

B

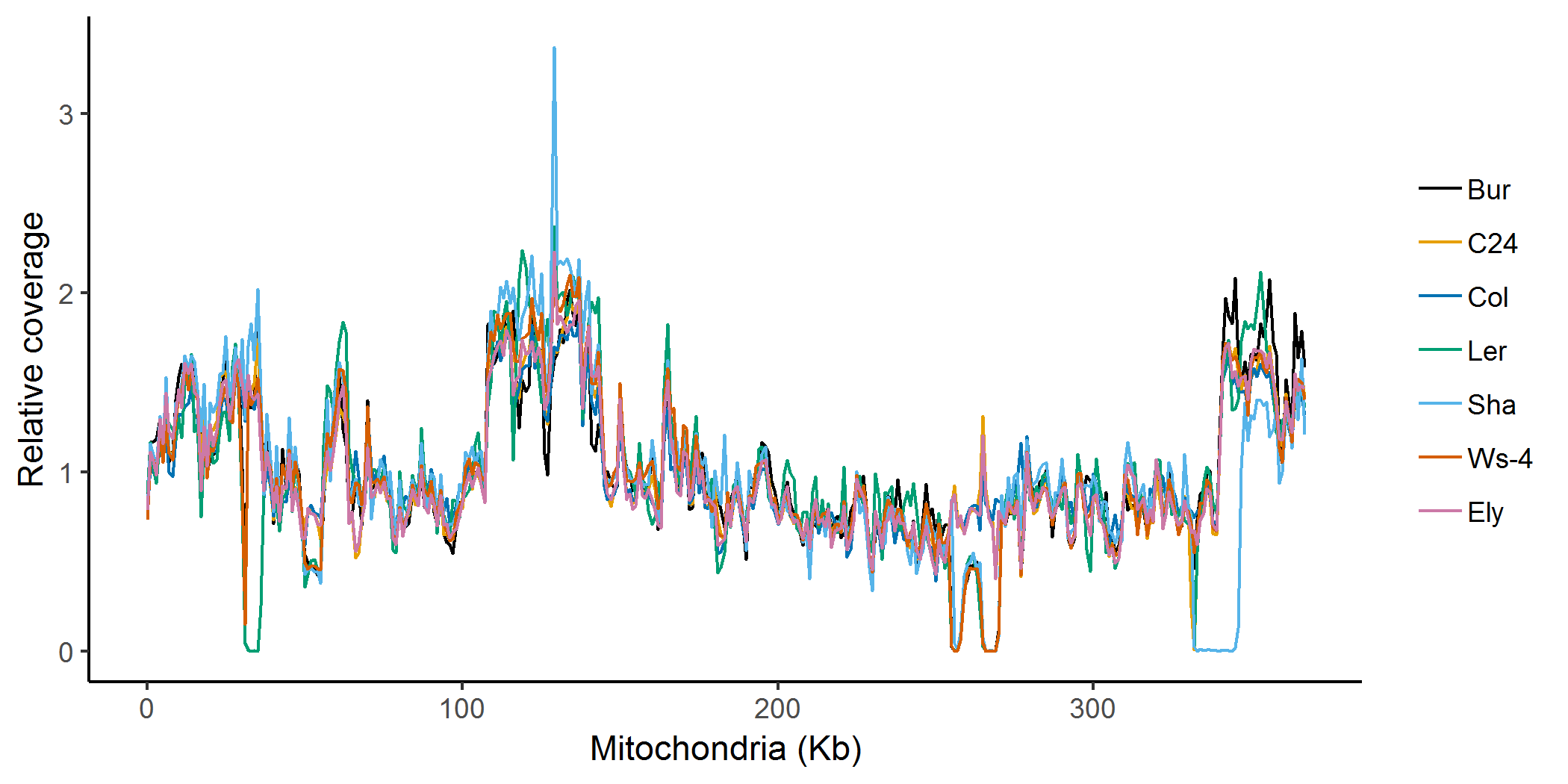

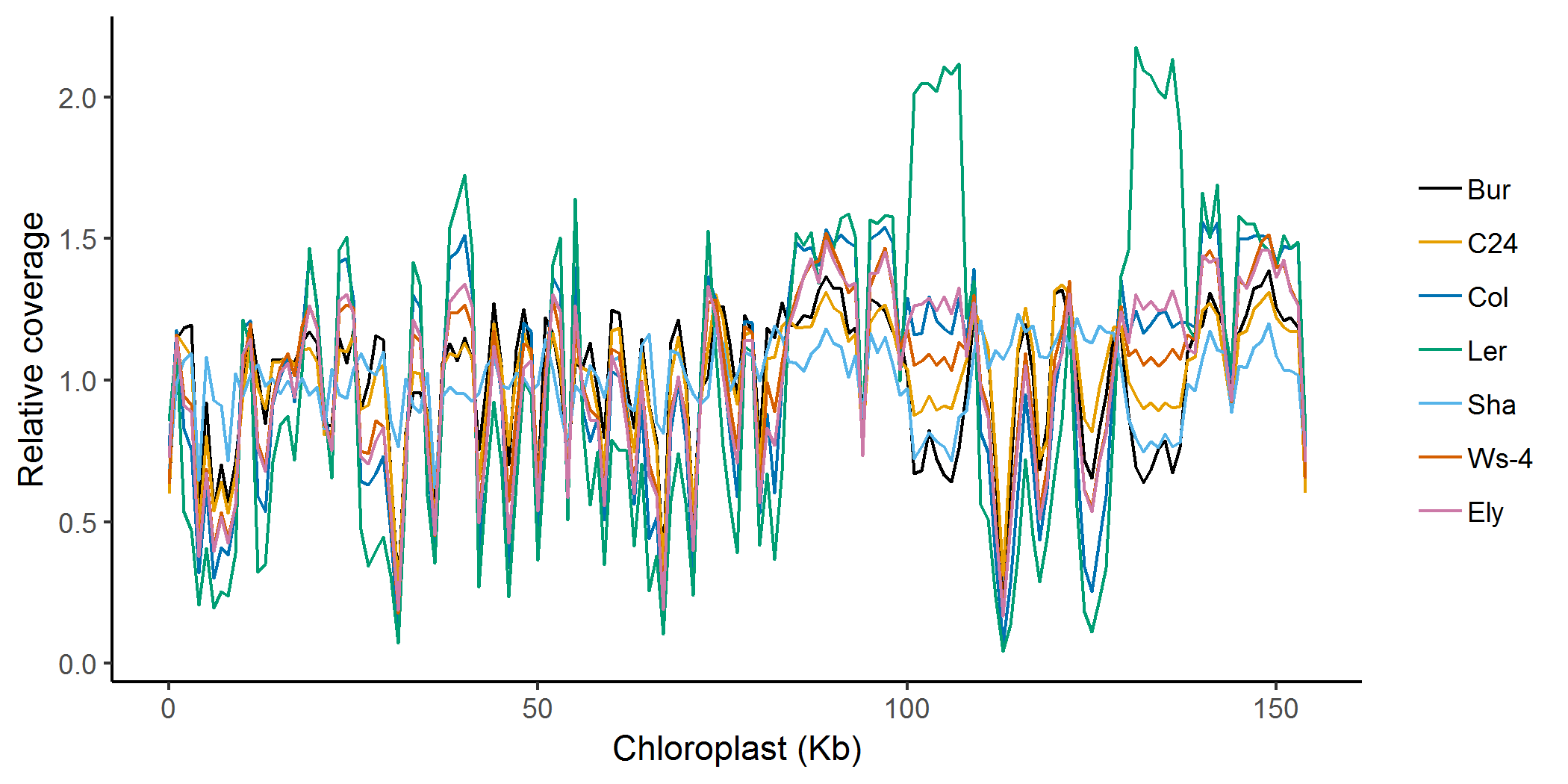

A

**Supplementary Figure 12.** Relative read depth for the chloroplast (A) and mitochondrial (B) genome sequences calculated in a sliding window of 1-kb. We observe no unique deletions or duplications in the Bur plasmotype that might be causal to the phenotypic effects observed in cybrids with the Bur plasmotype.

**Supplementary Table 1.** Assessment of statistical significance of phenotypic comparisons between wild types and corresponding self-cybrids (i.e. a wild-type nucleotype-plasmotype combination that went through a *cenh3*-induced haploid phase). P-values given for the null hypothesis that they are not different. The data is shown for a selection of 92 phenotypes (description given in Supplementary Table 4). As Bur^Bur^ was excluded, p-values are calculated on the basis of the other six wild-type vs self-cybrid comparisons. Measurements on a certain day are indicated with “d” and measurements on a certain hour are indicated with “h”. The p-value for “Flowering time” is below the 0.05 threshold, but in the posthoc test none of the 6 wild-type vs self-cybrid comparsons shows a significant difference, which occasionally happens in ANOVA tests, and thus is no reason to assume a difference.

| **Phentoype** | **p-value** | **Phentoype** | **p-value** |
| --- | --- | --- | --- |
| phiPSII (DEPI Exp 1) - 17.96h | 0.8113 | CD 2.5days - t50 maxG (Cs, hr) | 0.8780 |
| phiPSII (DEPI Exp 1) - 38.46h | 0.6714 | CD 5days - t50 maxG (Cs, hr) | 0.8095 |
| phiPSII (DEPI Exp 1) - 41.96h | 0.6300 | CD 7days - t50 maxG (Cs, hr) | 0.9063 |
| phiPSII (DEPI Exp 1) - 66.79h | 0.3721 | H1 no strat - t50 maxG (Cs, hr) | 0.5875 |
| phiPSII (DEPI Exp 1) - 71.46h | 0.2278 | H1 strat - t50 maxG (Cs, hr) | 0.9881 |
| phiNPQ (DEPI Exp 1) - 17.96h | 0.8722 | H2 strat - t50 maxG (Cs, hr) | 0.1936 |
| phiNPQ (DEPI Exp 1) - 38.46h | 0.7247 | NaCl - t50 maxG (Cs, hr) | 0.2640 |
| phiNPQ (DEPI Exp 1) - 41.96h | 0.7555 | Flowering time - First Flower | 0.0258 |
| phiNPQ (DEPI Exp 1) - 66.79h | 0.5000 | Seed Size - Area | 0.5428 |
| phiNPQ (DEPI Exp 1) - 71.46h | 0.5476 | Pollen Abortion | 0.7717 |
| phiNO (DEPI Exp 1) - 17.96h | 0.6402 | Metabolites - Boric acid (3TMS) | 0.1790 |
| phiNO (DEPI Exp 1) - 38.46h | 0.6779 | Metabolites - Benzeneacetonitrile, 2-cyano | 0.4985 |
| phiNO (DEPI Exp 1) - 41.96h | 0.5919 | Metabolites - Centrotype 670 | 0.0538 |
| phiNO (DEPI Exp 1) - 66.79h | 0.3803 | Metabolites - Serine (2TMS) | 0.5774 |
| phiNO (DEPI Exp 1) - 71.46h | 0.2205 | Metabolites - Ethanolamine (3TMS) | 0.0922 |
| NPQ (DEPI Exp 1) - 17.96h | 0.7688 | Metabolites - Glycerol (3TMS) | 0.0898 |
| NPQ (DEPI Exp 1) - 38.46h | 0.8983 | Metabolites - Threonine (2TMS) | 0.3861 |
| NPQ (DEPI Exp 1) - 41.96h | 0.9515 | Metabolites - Glycine (3TMS) | 0.9281 |
| NPQ (DEPI Exp 1) - 66.79h | 0.8783 | Metabolites - Glyceric acid (3TMS) | 0.2287 |
| NPQ (DEPI Exp 1) - 71.46h | 0.9278 | Metabolites - Fumaric acid (2TMS) | 0.9323 |
| QESV (DEPI Exp 1) - 17.96h | 0.7437 | Metabolites - Malic acid (3TMS) | 0.5128 |
| QESV (DEPI Exp 1) - 38.46h | 0.6961 | Metabolites - Pyroglutamic acid (2TMS) | 0.8601 |
| QESV (DEPI Exp 1) - 41.96h | 0.8290 | Metabolites - Glutamic acid (2TMS) | 0.2982 |
| QESV (DEPI Exp 1) - 66.79h | 0.1946 | Metabolites - Threonic acid (4TMS) | 0.9935 |
| QESV (DEPI Exp 1) - 71.46h | 0.2947 | Metabolites - 1,6-anhydro Glucose (3TMS) | 0.5133 |
| QI (DEPI Exp 1) - 17.99h | 0.9011 | Metabolites - Centrotype 3180 | 0.5933 |
| QI (DEPI Exp 1) - 38.49h | 0.8487 | Metabolites - Centrotype 3322 | 0.2838 |
| QI (DEPI Exp 1) - 41.99h | 0.8577 | Metabolites - Centrotype 3357 | 0.9074 |
| QI (DEPI Exp 1) - 66.83h | 0.4770 | Metabolites - Centrotype 3425 | 0.4037 |
| QI (DEPI Exp 1) - 71.49h | 0.5498 | Metabolites - Glucopyranose [-H20] (4TMS) | 0.4071 |
| FqFm (Phenovator) - Low Light 15.38d | 0.5357 | Metabolites - Centrotype 3532 | 0.3365 |
| FqFm (Phenovator) - Low Light 19.38d | 0.1924 | Metabolites - Dimethyl(trimethylsilyl)methoxysilane | 0.7407 |
| FqFm (Phenovator) - High Light 23.38d | 0.5113 | Metabolites - Centrotype 3597 | 0.1532 |
| FqFm (Phenovator) – Ratio | 0.5185 | Metabolites - Arabinofuranose (4TMS) | 0.4793 |
| Leaf Movement (Phenovator) - 16.54d | 0.6196 | Metabolites - Citric acid (4TMS) | 0.9022 |
| PLA (Phenovator) - 15.38d | 0.8623 | Metabolites - Centrotype 3795 | 0.5743 |
| PLA (Phenovator) - 19.38d | 0.5141 | Metabolites - Ascorbic acid (4TMS) | 0.3633 |
| PLA (Phenovator) - 23.38d | 0.8435 | Metabolites - Sorbose (5TMS) | 0.3672 |
| Chlorophyll Content (Phenovator) - 15.82d | 0.0790 | Metabolites - Fructose, methyloxime, (syn) (5TMS) | 0.3897 |
| Chlorophyll Content (Phenovator) - 23.82d | 0.1260 | Metabolites - Mannose, methyloxyme, (1Z) (5TMS) | 0.2884 |
| Biomass (D2) - 30d | 0.4719 | Metabolites - Centrotype 4295 | 0.3119 |
| PLA (D2 ) - 8.46d | 0.2468 | Metabolites - Glucose,methyloxime, (1E) (5TMS) | 0.4106 |
| PLA (D2 ) - 13.42d | 0.3214 | Metabolites - Glucose,methyloxime, (1Z) (5TMS) | 0.4976 |
| PLA (D2 ) - 18.42d | 0.0743 | Metabolites - 3,6-Dioxa-2,7-disilaoctane, 2,2,4,7,7-pentamethyl | 0.8775 |
| PLA (D2 ) - 23.42d | 0.1635 | Metabolites - Centrotype 5125 | 0.9815 |
| PLA Relative (D2 ) - Median rgr | 0.5393 | Metabolites - Centrotype 5879 | 0.2001 |

**Supplementary Table 2.** Contributions of nucleotype, plasmotype and their epistatic interaction (“Nucleotype:Plasmotype”) to the broad sense heritability (H^2^) for the selected subset of 92 phenotypes, excluding the Ely plasmotype. Measurements on a certain day are indicated with “d” and measurements on a certain hour are indicated with “h”.

| **Phentoype** | **Nucleotype:Plasmotype** | **Plasmotype** | **Nucleotype** |
| --- | --- | --- | --- |
| phiPSII (DEPI Exp 1) - 17.96h | 2.20% | 2.54% | 95.26% |
| phiPSII (DEPI Exp 1) - 38.46h | 1.42% | 0.79% | 97.80% |
| phiPSII (DEPI Exp 1) - 41.96h | 0.00% | 0.43% | 99.57% |
| phiPSII (DEPI Exp 1) - 66.79h | 0.00% | 1.04% | 98.96% |
| phiPSII (DEPI Exp 1) - 71.46h | 2.18% | 6.94% | 90.89% |
| phiNPQ (DEPI Exp 1) - 17.96h | 0.00% | 0.00% | 100.00% |
| phiNPQ (DEPI Exp 1) - 38.46h | 9.35% | 0.00% | 90.65% |
| phiNPQ (DEPI Exp 1) - 41.96h | 0.00% | 0.00% | 100.00% |
| phiNPQ (DEPI Exp 1) - 66.79h | 2.28% | 0.00% | 97.72% |
| phiNPQ (DEPI Exp 1) - 71.46h | 17.82% | 7.54% | 74.64% |
| phiNO (DEPI Exp 1) - 17.96h | 0.80% | 2.07% | 97.13% |
| phiNO (DEPI Exp 1) - 38.46h | 0.00% | 3.31% | 96.69% |
| phiNO (DEPI Exp 1) - 41.96h | 0.02% | 4.22% | 95.76% |
| phiNO (DEPI Exp 1) - 66.79h | 0.00% | 2.65% | 97.35% |
| phiNO (DEPI Exp 1) - 71.46h | 0.97% | 4.56% | 94.48% |
| NPQ (DEPI Exp 1) - 17.96h | 0.00% | 0.09% | 99.91% |
| NPQ (DEPI Exp 1) - 38.46h | 5.99% | 3.51% | 90.50% |
| NPQ (DEPI Exp 1) - 41.96h | 0.00% | 2.36% | 97.64% |
| NPQ (DEPI Exp 1) - 66.79h | 3.89% | 0.66% | 95.45% |
| NPQ (DEPI Exp 1) - 71.46h | 2.95% | 0.60% | 96.45% |
| QESV (DEPI Exp 1) - 17.96h | 3.17% | 0.00% | 96.83% |
| QESV (DEPI Exp 1) - 38.46h | 5.85% | 0.00% | 94.15% |
| QESV (DEPI Exp 1) - 41.96h | 0.00% | 0.00% | 100.00% |
| QESV (DEPI Exp 1) - 66.79h | 0.47% | 0.00% | 99.53% |
| QESV (DEPI Exp 1) - 71.46h | 5.83% | 21.99% | 72.18% |
| QI (DEPI Exp 1) - 17.99h | 0.00% | 0.51% | 99.49% |
| QI (DEPI Exp 1) - 38.49h | 0.77% | 4.43% | 94.80% |
| QI (DEPI Exp 1) - 41.99h | 0.18% | 2.04% | 97.78% |
| QI (DEPI Exp 1) - 66.83h | 1.00% | 2.66% | 96.34% |
| QI (DEPI Exp 1) - 71.49h | 2.17% | 4.61% | 93.21% |
| FqFm (Phenovator) - Low Light 15.38d | 0.68% | 0.08% | 99.24% |
| FqFm (Phenovator) - Low Light 19.38d | 0.43% | 1.13% | 98.44% |
| FqFm (Phenovator) - High Light 23.38d | 0.00% | 0.04% | 99.96% |
| FqFm (Phenovator) – Ratio | 4.29% | 1.96% | 93.75% |
| Leaf Movement (Phenovator) - 16.54d | 8.27% | 0.39% | 91.34% |
| PLA (Phenovator) - 15.38d | 5.03% | 1.09% | 93.89% |
| PLA (Phenovator) - 19.38d | 4.05% | 0.88% | 95.07% |
| PLA (Phenovator) - 23.38d | 0.79% | 0.56% | 98.65% |
| Chlorophyll Content (Phenovator) - 15.82d | 0.00% | 0.72% | 99.28% |
| Chlorophyll Content (Phenovator) - 23.82d | 1.09% | 1.43% | 97.49% |
| Biomass (D2) - 30d | 7.37% | 2.61% | 90.02% |
| PLA (D2 ) - 8.46d | 9.34% | 0.00% | 90.66% |
| PLA (D2 ) - 13.42d | 12.27% | 0.00% | 87.73% |
| PLA (D2 ) - 18.42d | 12.30% | 0.00% | 87.70% |
| PLA (D2 ) - 23.42d | 7.99% | 0.00% | 92.01% |
| PLA Relative (D2 ) - Median rgr | 1.43% | 0.00% | 98.57% |
| CD 2.5days - t50 maxG (Cs, hr) | 4.27% | 0.14% | 95.59% |
| CD 5days - t50 maxG (Cs, hr) | 2.96% | 0.00% | 97.04% |
| CD 7days - t50 maxG (Cs, hr) | 3.69% | 0.00% | 96.31% |
| H1 no strat - t50 maxG (Cs, hr) | 6.96% | 0.00% | 93.04% |
| H1 strat - t50 maxG (Cs, hr) | 4.47% | 1.65% | 93.88% |
| H2 strat - t50 maxG (Cs, hr) | 6.57% | 0.00% | 93.43% |
| NaCl - t50 maxG (Cs, hr) | 5.80% | 0.00% | 94.20% |
| Flowering - First Flower | 2.32% | 0.35% | 97.33% |
| Seed Size – Area | 5.26% | 0.24% | 94.50% |
| Pollen Abortion | 43.49% | 1.39% | 55.11% |
| Metabolites - Boric acid (3TMS) | 34.59% | 0.00% | 65.41% |
| Metabolites - Benzeneacetonitrile, 2-cyano | 0.00% | 0.50% | 99.50% |
| Metabolites - Centrotype 670 | 1.76% | 0.00% | 98.24% |
| Metabolites - Serine (2TMS) | 0.00% | 5.41% | 94.59% |
| Metabolites - Ethanolamine (3TMS) | 23.33% | 11.66% | 65.01% |
| Metabolites - Glycerol (3TMS) | 0.00% | 0.00% | 100.00% |
| Metabolites - Threonine (2TMS) | 0.00% | 3.32% | 96.68% |
| Metabolites - Glycine (3TMS) | 0.00% | 8.32% | 91.68% |
| Metabolites - Glyceric acid (3TMS) | 0.00% | 19.34% | 80.66% |
| Metabolites - Fumaric acid (2TMS) | 0.00% | 0.00% | 100.00% |
| Metabolites - Malic acid (3TMS) | 0.00% | 0.00% | 100.00% |
| Metabolites - Pyroglutamic acid (2TMS) | 0.00% | 0.00% | 100.00% |
| Metabolites - Glutamic acid (2TMS) | 0.00% | 0.00% | 100.00% |
| Metabolites - Threonic acid (4TMS) | 0.00% | 0.00% | 100.00% |
| Metabolites - 1,6-anhydro Glucose (3TMS) | 0.00% | 0.00% | 100.00% |
| Metabolites - Centrotype 3180 | 0.00% | 13.15% | 86.85% |
| Metabolites - Centrotype 3322 | 0.00% | 0.00% | 100.00% |
| Metabolites - Centrotype 3357 | 18.61% | 18.41% | 62.98% |
| Metabolites - Centrotype 3425 | 0.00% | 0.00% | 100.00% |
| Metabolites - Glucopyranose [-H20] (4TMS) | 0.00% | 0.00% | 100.00% |
| Metabolites - Centrotype 3532 | 10.83% | 2.31% | 86.86% |
| Metabolites - Dimethyl(trimethylsilyl)methoxysilane | 0.00% | 0.00% | 100.00% |
| Metabolites - Centrotype 3597 | 0.00% | 0.00% | 100.00% |
| Metabolites - Arabinofuranose (4TMS) | 10.59% | 2.88% | 86.53% |
| Metabolites - Citric acid (4TMS) | 20.85% | 0.00% | 79.15% |
| Metabolites - Centrotype 3795 | 11.54% | 0.00% | 88.46% |
| Metabolites - Ascorbic acid (4TMS) | 0.00% | 1.41% | 98.59% |
| Metabolites - Sorbose (5TMS) | 0.00% | 1.60% | 98.40% |
| Metabolites - Fructose, methyloxime, (syn) (5TMS) | 0.00% | 0.00% | 100.00% |
| Metabolites - Mannose, methyloxyme, (1Z) (5TMS) | 59.82% | 0.00% | 40.18% |
| Metabolites - Centrotype 4295 | 0.00% | 1.68% | 98.32% |
| Metabolites - Glucose,methyloxime, (1E) (5TMS) | 0.00% | 2.77% | 97.23% |
| Metabolites - Glucose,methyloxime, (1Z) (5TMS) | 52.63% | 47.37% | 0.00% |
| Metabolites - 3,6-Dioxa-2,7-disilaoctane, 2,2,4,7,7-pentamethyl | 0.00% | 34.39% | 65.61% |
| Metabolites - Centrotype 5125 | 0.00% | 0.00% | 100.00% |
| Metabolites - Centrotype 5879 | 0.00% | 1.55% | 98.45% |

**Supplementary Table 3.** Average and standard deviation for the contribution of genetic components to the broad sense heritability (H^2^). In the upper table, the Ely plasmotype is excluded to remove its known additive effect, which facilitates the exploration of the remaining contribution of the plasmotype. In the lower table the Ely plasmotype is included. Data given for 1782 phenotypes (1859, excluding the 77 phenotypes with H^2^ lower than 5%) together, the selected subset of 92 phenotypes. Percentage explained genetic variance is calculated for the three genetic components (nucleotype, plasmotype and nucleotype-plasmotype interaction). See Supplementary Data 1 for the complete dataset.

*Without the Ely plasmotype*

|  | **Genetic Component** | **Mean** | **Standard Deviation** |
| --- | --- | --- | --- |
| **All 1782 Phenotypes** | Nucleotype | 94.89% | 8.19% |
|  | Nucleotype:Plasmotype | 2.33% | 5.38% |
|  | Plasmotype | 2.78% | 5.99% |
| **Subset of 92 phenotypes** | Nucleotype | 91.90% | 14.40% |
|  | Nucleotype:Plasmotype | 5.16% | 10.48% |
|  | Plasmotype | 2.94% | 7.03% |

*With the Ely plasmotype*

|  | **Genetic Component** | **Mean** | **Standard Deviation** |
| --- | --- | --- | --- |
| **All 1782 Phenotypes** | Nucleotype | 45.71% | 30.21% |
|  | Nucleotype:Plasmotype | 6.30% | 8.66% |
|  | Plasmotype | 47.99% | 33.76% |
| **Subset of 92 phenotypes** | Nucleotype | 65.90% | 29.32% |
|  | Nucleotype:Plasmotype | 6.08% | 9.51% |
|  | Plasmotype | 28.02% | 29.99% |

**Supplementary Table 4.** Overview of all 1859 phenotypes for 17 traits used in this study. A small discription is given for every trait. The phenotyping platform for every trait is mentioned, as described in the methods. The statistical model used together with the input data, raw data means unadjusted data, fitted data means smoothend by fitting a spline and batch correction refers to batch correction as described in the methods. See Supplementary Data 1 for complete list of phenotypes.

| **Trait** | **Abbreviation** | **Description of trait** | **Phenotyping Platform** | **Number of phenotypes** | **Replicate number (n)** | **Form of data used** | **Statistical Model** |
| --- | --- | --- | --- | --- | --- | --- | --- |
| PSII Efficiency | Φpsii, Fq/Fm | PSII efficiency for electron transport | DEPI | 84 + 144 | 4 + 4 | Raw Data | $\underline{Y}= {Nucleotype}+Plasmotype+(Nucleotype*Plasmotype)+ \underline{\varepsilon}$ |
| phiNO | ΦNO | Basal dissipation | DEPI | 84 + 144 | 4 + 4 | Raw Data | $\underline{Y}= {Nucleotype}+Plasmotype+(Nucleotype*Plasmotype)+ \underline{\varepsilon}$ |
| phiNPQ | ΦNPQ | Inducible dissipation | DEPI | 84 + 144 | 4 + 4 | Raw Data | $\underline{Y}= {Nucleotype}+Plasmotype+(Nucleotype*Plasmotype)+ \underline{\varepsilon}$ |
| NPQ | - | Non-photochemical quenching | DEPI | 84 + 144 | 4 + 4 | Raw Data | $\underline{Y}= {Nucleotype}+Plasmotype+(Nucleotype*Plasmotype)+ \underline{\varepsilon}$ |
| QESV | q_E_ | Rapidly relaxing component of NPQ | DEPI | 83 + 144 | 4 + 4 | Raw Data | $\underline{Y}= {Nucleotype}+Plasmotype+(Nucleotype*Plasmotype)+ \underline{\varepsilon}$ |
| QI | q_I_ | Slowly relaxing component of NPQ | DEPI | 83 + 144 | 4 + 4 | Raw Data | $\underline{Y}= {Nucleotype}+Plasmotype+(Nucleotype*Plasmotype)+ \underline{\varepsilon}$ |
| PSII Efficiency | Φpsii, Fq/Fm | PSII efficiency for electron transport | Phenovator | 40 | 24 | Raw Data | $\underline{Y}= {Nucleotype}+Plasmotype+(Nucleotype*Plasmotype)+\underline{Basin}+ \underline{Table Position}+\underline{Image Position}+ \underline{\varepsilon}$ |
| Plant leaf area | PLA | Absolute plant leaf area | Phenovator | 51 | 24 | Raw Data | $\underline{Y}= {Nucleotype}+Plasmotype+(Nucleotype*Plasmotype)+\underline{Basin}+ \underline{Table Position}+\underline{Image Position}+ \underline{\varepsilon}$ |
| Chlorophyll Reflectance | - | Reflectance-based estimate of chlorophyll | Phenovator | 13 | 24 | Raw Data | $\underline{Y}= {Nucleotype}+Plasmotype+(Nucleotype*Plasmotype)+\underline{Basin}+ \underline{Table Position}+\underline{Image Position}+ \underline{\varepsilon}$ |
| Leaf Movement | - | Epinastic leaf movement, relative to the previous measurement | Phenovator | 99 | 24 | Raw Data | $\underline{Y}= {Nucleotype}+Plasmotype+(Nucleotype*Plasmotype)+\underline{Basin}+ \underline{Table Position}+\underline{Image Position}+ \underline{\varepsilon}$ |
| Growth | - | Absolute plant leaf area | D2 | 56 | 24 | Raw Data | $\underline{Y}= {Nucleotype}+Plasmotype+(Nucleotype*Plasmotype)+ \underline{\varepsilon}$ |
|  | Fitted | Fitted plant leaf area | D2 | 64 | 24 | Fitted Data | $\underline{Y}= {Nucleotype}+Plasmotype+(Nucleotype*Plasmotype)+\underline{Block}+ \underline{\varepsilon}$ |
|  | RGR | Relative Growth Rate | D2 | 62 | 24 | Fitted Data | $\underline{Y}= {Nucleotype}+Plasmotype+(Nucleotype*Plasmotype)+\underline{Block}+ \underline{\varepsilon}$ |
| Biomass | - | Above ground shoot dry weight (g) | D2 | 1 | 12 | Raw Data | $\underline{Y}= {Nucleotype}+Plasmotype+(Nucleotype*Plasmotype)+\underline{Block}+ \underline{\varepsilon}$ |
| Germination | - |  | Germinator | 63 | 4 | Raw Data | $\underline{Y}= {Nucleotype}+Plasmotype+(Nucleotype*Plasmotype)+ \underline{\varepsilon}$ |
| Seed size | - |  | Germinator | 2 | 10 | Raw Data | $\underline{Y}= {Nucleotype}+Plasmotype+(Nucleotype*Plasmotype)+ \underline{\varepsilon}$ |
| Flowering Time | Ft | Days after sowing for the first flower to open | Scored by hand | 2 | 10 | Raw Data | $\underline{Y}= {Nucleotype}+Plasmotype+(Nucleotype*Plasmotype)+\underline{Block}+ \underline{\varepsilon}$ |
| Pollen abortion | - | # of aborted pollen out of 250 | Counted with microscope | 1 | 4-10 | Raw Data | $\underline{Y}= {Nucleotype}+Plasmotype+(Nucleotype*Plasmotype)+ \underline{\varepsilon}$ |
| Metabolites | - | Metabolite concentration in above ground shoot | GC-ToF-MS | 39 | 4 (pools of 6 plants) | Batch Corrected Data | $\underline{Y}= {Nucleotype}+Plasmotype+(Nucleotype*Plasmotype)+ \underline{\varepsilon}$ |

| **Plasmotype Comparison** | **Nucleotype** | **Total # detected genes** | **DE (α=0.05)** | **Percentage DE (%)** | **Up-regulated** | **Down-regulated** | **Percentage down compared to DE** |
| --- | --- | --- | --- | --- | --- | --- | --- |
| Bur vs L*er* | L*er* | 22376 | 3 | 0.01% | 1 | 2 | 66.7% |
| Bur vs L*er* | Ely | 20205 | 117 | 0.58% | 112 | 5 | 4.3% |
| Ely vs Bur | L*er* | 17130 | 458 | 2.67% | 267 | 191 | 41.7% |
| Ely vs Bur | Ely | 17929 | 1293 | 7.21% | 845 | 448 | 34.6% |
| Ely vs L*er* | L*er* | 15410 | 726 | 4.71% | 426 | 300 | 41.3% |
| Ely vs L*er* | Ely | 5897 | 189 | 3.21% | 181 | 8 | 4.2% |

**Supplementary Table 5.** Gene expression counts in pairwise comparisons of cybrids with L*er* or Ely nucleotypes and Bur, L*er* or Ely plasmotypes. For each comparison the total number of detected genes, and the number of differentially expressed (DE) genes, both up- and down-regulated, is given. P-value correction using the Hochberg correction (α=0.05), n=6.

**Supplementary Table 6.** Genes that are become differentially expressed due to the Ely plasmotype. When in an Ely or L*er* nuclear background the Ely plasmotype is replaced by L*er* or Bur, the below listed genes are significantly differentially expressed. These genes can be interpreted as a main effect of replacing the Ely plasmotype by a Bur or L*er* plasmotype. All genes (except AT3G57240) become upregulated when the Ely plasmotype is replaced.

| **Gene** | **Name** | **Description** |
| --- | --- | --- |
| AT1G11860 | AT1G11860 | Glycine cleavage T-protein family |
| AT1G31800 | CYP97A3 | Protein LUTEIN DEFICIENT 5, chloroplastic |
| AT1G42970 | GAPB | Glyceraldehyde-3-phosphate dehydrogenase GAPB, chloroplastic |
| AT1G49380 | CCS1 | Cytochrome c biogenesis protein CCS1, chloroplastic |
| AT1G57770 | AT1G57770 | FAD/NAD(P)-binding oxidoreductase family protein |
| AT1G62750 | CPEFG | Elongation factor G, chloroplastic |
| AT1G64680 | AT1G64680 |  |
| AT1G68010 | HPR | hydroxypyruvate reductase |
| AT1G69523 | AT1G69523 |  |
| AT1G72030 | AT1G72030 | Acyl-CoA N-acyltransferases (NAT) superfamily protein |
| AT1G73325 | AT1G73325 | Kunitz family trypsin and protease inhibitor protein |
| AT2G13360 | AGT1 | Serine--glyoxylate aminotransferase |
| AT2G22360 | DJA6 | Chaperone protein dnaJ A6, chloroplastic |
| AT2G26080 | GLDP2 | Glycine dehydrogenase (decarboxylating) 2, mitochondrial |
| AT2G32500 | AT2G32500 |  |
| AT2G36145 | AT2G36145 | Expressed protein |
| AT2G38230 | PDX11 | Pyridoxal 5'-phosphate synthase subunit PDX1.1 |
| AT3G01500 | BCA1 | Beta carbonic anhydrase 1, chloroplastic |
| AT3G10840 | AT3G10840 | Alpha/beta-Hydrolases superfamily protein |
| AT3G14415 | AT3G14415 | GOX2 |
| AT3G14420 | GLO1 | (S)-2-hydroxy-acid oxidase GLO1 |
| AT3G19480 | PGDH3 | D-3-phosphoglycerate dehydrogenase 3, chloroplastic |
| AT3G48420 | CBBY | CBBY-like protein |
| AT3G55800 | SBPASE | Sedoheptulose-1,7-bisphosphatase, chloroplastic |
| AT3G57240 | BG3 | Probable glucan endo-1,3-beta-glucosidase BG3 |
| AT4G10120 | SPS4 | Probable sucrose-phosphate synthase 4 |
| AT4G12320 | CYP706A6 |  |
| AT4G12830 | AT4G12830 |  |
| AT4G27820 | BGLU9 | Beta-glucosidase 9 |
| AT4G28740 | AT4G28740 | FUNCTIONS IN: molecular_function unknown; LOCATED IN: chloroplast; |
| AT4G34090 | AT4G34090 | unknown protein; LOCATED IN: chloroplast, chloroplast stroma |
| AT4G36530 | AT4G36530 | Alpha/beta-Hydrolases superfamily protein |
| AT5G05580 | FAD8 | Temperature-sensitive sn-2 acyl-lipid omega-3 desaturase (ferredoxin), chloroplastic |
| AT5G09660 | PMDH2 | Malate dehydrogenase |
| AT5G14740 | BCA2 | Beta carbonic anhydrase 2, chloroplastic |
| AT5G38430 | RBCS-1B | Ribulose bisphosphate carboxylase small chain 1B, chloroplastic |
| AT5G51820 | PGMP | Phosphoglucomutase, chloroplastic |
| AT5G55740 | CRR21 | Pentatricopeptide repeat-containing protein At5g55740, chloroplastic |
| AT5G58260 | ndhN | NdhN |
| AT5G58310 | MES18 | Methylesterase 18 |

**Supplementary Table 7.** List of primers used for KASP analysis. Primer names show the plasmotype and the SNP position they detect (base pair position according to TAIR10 reference genome). A1 and A2 are forward primers, and C1 is the reverse primer.

| Primer name | Oligo sequence |
| --- | --- |
| Bur_15124_A1 | GAAGGTGACCAAGTTCATGCTACAAATTGTAATCCCGATCTCGCG |
| Bur_15124_A2 | GAAGGTCGGAGTCAACGGATTATACAAATTGTAATCCCGATCTCGCA |
| Bur_15124_C1 | GCTATAGCGTCATCATTTGCGGGAA |
| C24_11907_A1 | GAAGGTGACCAAGTTCATGCTGATTTATTAGATAACCGAAAGCAGAGG |
| C24_11907_A2 | GAAGGTCGGAGTCAACGGATTAATGATTTATTAGATAACCGAAAGCAGAGA |
| C24_11907_C1 | GCTCCTTCACGCAGTTCTTCTGAAT |
| Col_23915_A1 | GAAGGTGACCAAGTTCATGCTCACGTTTCTGTGAAATATATACACGAATT |
| Col_23915_A2 | GAAGGTCGGAGTCAACGGATTCACGTTTCTGTGAAATATATACACGAATG |
| Col_23915_C1 | GGTTCAGAAAAAGGGTGGTTCAAGTTATA |
| Ely_655_A1 | GAAGGTGACCAAGTTCATGCTTGGCCGATTGATTTTCCAATATGCTA |
| Ely_655_A2 | GAAGGTCGGAGTCAACGGATTGGCCGATTGATTTTCCAATATGCTG |
| Ely_655_C1 | GGCCAAGCCGCTAAGAAGAAATGTA |
| Ler_74355_A1 | GAAGGTGACCAAGTTCATGCTGAGGCTGATTACGTTAACTAGTCC |
| Ler_74355_A2 | GAAGGTCGGAGTCAACGGATTGGAGGCTGATTACGTTAACTAGTCT |
| Ler_74355_C1 | GGCAACCCTCTCAACAACTAAGAGAT |
| Sha_4975_A1 | GAAGGTGACCAAGTTCATGCTATCTACTCTTCTCTTTCACTTCCATCA |
| Sha_4975_A2 | GAAGGTCGGAGTCAACGGATTCTACTCTTCTCTTTCACTTCCATCG |
| Sha_4975_C1 | CCTTAGGAGGAATACTAATAATAAATAGAA |
| Ws-4_20000_A1 | GAAGGTGACCAAGTTCATGCTCCCGAAGTGATCTATTAATCGGCTA |
| Ws-4_20000_A2 | GAAGGTCGGAGTCAACGGATTCCGAAGTGATCTATTAATCGGCTG |
| Ws-4_20000_C1 | CACAATAAAGTGATAGATGGAACTGCTATT |

**Supplementary Table 8.** Outcome of the KASP^TM^ assay to check for the heterozygous calls in C24^C24^ and Ws-4^Col^. A KASP^TM^ assay requires two different forward primers, that only attach on the presence of a specific SNP, as such a KASP^TM^ assay can distinguish between being homozygous for allele 1 (TAIR 10 Col reference), allele 2 (plasmotype alternative, indicated on top row) or heterozygous. All 7 marker sets have been assayed on the complete panel (excluding Ely^Sha^). The markers were designed for unique SNPs, the combination of the 7 markers sets represents a unique chloroplast genome. From the absence of heterozygous calls, we conclude that all cybrids are correct including C24^C24^ and Ws-4^Col^. Blue indicates whether the cybrids contains allele 1 and orange indicates whether it contains allele 2.

| **Nucleotype** | **Plasmotype** | **Bur** | **C24** | **Col** | **Ely** | **Ler** | **Sha** | **Ws-4** |
| --- | --- | --- | --- | --- | --- | --- | --- | --- |
| WT | Bur-0 | Allele 2 | Allele 1 | Allele 1 | Allele 1 | Allele 1 | Allele 1 | Allele 1 |
| Bur-0 | Bur-0 | Allele 2 | Allele 1 | Allele 1 | Allele 1 | Allele 1 | Allele 1 | Allele 1 |
| C24 | Bur-0 | Allele 2 | Allele 1 | Allele 1 | Allele 1 | Allele 1 | Allele 1 | Allele 1 |
| Col-0 | Bur-0 | Allele 2 | Allele 1 | Allele 1 | Allele 1 | Allele 1 | Allele 1 | Allele 1 |
| Ely | Bur-0 | Allele 2 | Allele 1 | Allele 1 | Allele 1 | Allele 1 | Allele 1 | Allele 1 |
| Ler-1 | Bur-0 | Allele 2 | Allele 1 | Allele 1 | Allele 1 | Allele 1 | Allele 1 | Allele 1 |
| Shah | Bur-0 | Allele 2 | Allele 1 | Allele 1 | Allele 1 | Allele 1 | Allele 1 | Allele 1 |
| Ws-4 | Bur-0 | Allele 2 | Allele 1 | Allele 1 | Allele 1 | Allele 1 | Allele 1 | Allele 1 |
| WT | C24 | Allele 1 | Allele 2 | Allele 1 | Allele 1 | Allele 1 | Allele 1 | Allele 1 |
| Bur-0 | C24 | Allele 1 | Allele 2 | Allele 1 | Allele 1 | Allele 1 | Allele 1 | Allele 1 |
| C24 | C24 | Allele 1 | Allele 2 | Allele 1 | Allele 1 | Allele 1 | Allele 1 | Allele 1 |
| Col-0 | C24 | Allele 1 | Allele 2 | Allele 1 | Allele 1 | Allele 1 | Allele 1 | Allele 1 |
| Ely | C24 | Allele 1 | Allele 2 | Allele 1 | Allele 1 | Allele 1 | Allele 1 | Allele 1 |
| Ler-1 | C24 | Allele 1 | Allele 2 | Allele 1 | Allele 1 | Allele 1 | Allele 1 | Allele 1 |
| Shah | C24 | Allele 1 | Allele 2 | Allele 1 | Allele 1 | Allele 1 | Allele 1 | Allele 1 |
| Ws-4 | C24 | Allele 1 | Allele 2 | Allele 1 | Allele 1 | Allele 1 | Allele 1 | Allele 1 |
| WT | Col-0 | Allele 2 | Allele 2 | Allele 1 | Allele 2 | Allele 2 | Allele 2 | Allele 2 |
| Bur-0 | Col-0 | Allele 2 | Allele 2 | Allele 1 | Allele 2 | Allele 2 | Allele 2 | Allele 2 |
| C24 | Col-0 | Allele 2 | Allele 2 | Allele 1 | Allele 2 | Allele 2 | Allele 2 | Allele 2 |
| Col-0 | Col-0 | Allele 2 | Allele 2 | Allele 1 | Allele 2 | Allele 2 | Allele 2 | Allele 2 |
| Ely | Col-0 | Allele 2 | Allele 2 | Allele 1 | Allele 2 | Allele 2 | Allele 2 | Allele 2 |
| Ler-1 | Col-0 | Allele 2 | Allele 2 | Allele 1 | Allele 2 | Allele 2 | Allele 2 | Allele 2 |
| Shah | Col-0 | Allele 2 | Allele 2 | Allele 1 | Allele 2 | Allele 2 | Allele 2 | Allele 2 |
| Ws-4 | Col-0 | Allele 2 | Allele 2 | Allele 1 | Allele 2 | Allele 2 | Allele 2 | Allele 2 |
| WT | Ely | Allele 1 | Allele 1 | Allele 1 | Allele 2 | Allele 1 | Allele 1 | Allele 1 |
| Bur-0 | Ely | Allele 1 | Allele 1 | Allele 1 | Allele 2 | Allele 1 | Allele 1 | Allele 1 |
| C24 | Ely | Allele 1 | Allele 1 | Allele 1 | Allele 2 | Allele 1 | Allele 1 | Allele 1 |
| Col-0 | Ely | Allele 1 | Allele 1 | Allele 1 | Allele 2 | Allele 1 | Allele 1 | Allele 1 |
| Ely | Ely | Allele 1 | Allele 1 | Allele 1 | Allele 2 | Allele 1 | Allele 1 | Allele 1 |
| Ler-1 | Ely | Allele 1 | Allele 1 | Allele 1 | Allele 2 | Allele 1 | Allele 1 | Allele 1 |
| Shah | Ely | Allele 1 | Allele 1 | Allele 1 | Allele 2 | Allele 1 | Allele 1 | Allele 1 |
| Ws-4 | Ely | Allele 1 | Allele 1 | Allele 1 | Allele 2 | Allele 1 | Allele 1 | Allele 1 |
| WT | Ler-1 | Allele 1 | Allele 1 | Allele 1 | Allele 1 | Allele 2 | Allele 1 | Allele 1 |
| Bur-0 | Ler-1 | Allele 1 | Allele 1 | Allele 1 | Allele 1 | Allele 2 | Allele 1 | Allele 1 |
| C24 | Ler-1 | Allele 1 | Allele 1 | Allele 1 | Allele 1 | Allele 2 | Allele 1 | Allele 1 |
| Col-0 | Ler-1 | Allele 1 | Allele 1 | Allele 1 | Allele 1 | Allele 2 | Allele 1 | Allele 1 |
| Ely | Ler-1 | Allele 1 | Allele 1 | Allele 1 | Allele 1 | Allele 2 | Allele 1 | Allele 1 |
| Ler-1 | Ler-1 | Allele 1 | Allele 1 | Allele 1 | Allele 1 | Allele 2 | Allele 1 | Allele 1 |
| Shah | Ler-1 | Allele 1 | Allele 1 | Allele 1 | Allele 1 | Allele 2 | Allele 1 | Allele 1 |
| Ws-4 | Ler-1 | Allele 1 | Allele 1 | Allele 1 | Allele 1 | Allele 2 | Allele 1 | Allele 1 |
| WT | Shah | Allele 1 | Allele 1 | Allele 1 | Allele 1 | Allele 1 | Allele 2 | Allele 1 |
| Bur-0 | Shah | Allele 1 | Allele 1 | Allele 1 | Allele 1 | Allele 1 | Allele 2 | Allele 1 |
| C24 | Shah | Allele 1 | Allele 1 | Allele 1 | Allele 1 | Allele 1 | Allele 2 | Allele 1 |
| Col-0 | Shah | Allele 1 | Allele 1 | Allele 1 | Allele 1 | Allele 1 | Allele 2 | Allele 1 |
| Ely | Shah | nd | nd | nd | nd | nd | Allele 2 | nd |
| Ler-1 | Shah | Allele 1 | Allele 1 | Allele 1 | Allele 1 | Allele 1 | Allele 2 | Allele 1 |
| Shah | Shah | Allele 1 | Allele 1 | Allele 1 | Allele 1 | Allele 1 | Allele 2 | Allele 1 |
| Ws-4 | Shah | Allele 1 | Allele 1 | Allele 1 | Allele 1 | Allele 1 | Allele 2 | Allele 1 |
| WT | Ws-4 | Allele 1 | Allele 1 | Allele 1 | Allele 1 | Allele 1 | Allele 1 | Allele 2 |
| Bur-0 | Ws-4 | Allele 1 | Allele 1 | Allele 1 | Allele 1 | Allele 1 | Allele 1 | Allele 2 |
| C24 | Ws-4 | Allele 1 | Allele 1 | Allele 1 | Allele 1 | Allele 1 | Allele 1 | Allele 2 |
| Col-0 | Ws-4 | Allele 1 | Allele 1 | Allele 1 | Allele 1 | Allele 1 | Allele 1 | Allele 2 |
| Ely | Ws-4 | Allele 1 | Allele 1 | Allele 1 | Allele 1 | Allele 1 | Allele 1 | Allele 2 |
| Ler-1 | Ws-4 | Allele 1 | Allele 1 | Allele 1 | Allele 1 | Allele 1 | Allele 1 | Allele 2 |
| Shah | Ws-4 | Allele 1 | Allele 1 | Allele 1 | Allele 1 | Allele 1 | Allele 1 | Allele 2 |
| Ws-4 | Ws-4 | Allele 1 | Allele 1 | Allele 1 | Allele 1 | Allele 1 | Allele 1 | Allele 2 |
